# A global, integrated view of the ubiquitylation site occupancy and dynamics

**DOI:** 10.1101/2023.07.19.549470

**Authors:** Gabriela Prus, Shankha Satpathy, Brian T. Weinert, Takeo Narita, Chunaram Choudhary

## Abstract

Ubiquitylation regulates virtually all proteins and biological processes in a cell. However, the global site-specific occupancy (stoichiometry) and turnover rate of ubiquitylation have never been quantified. Here, we present the first integrated picture of ubiquitylation site occupancy and half-life. Ubiquitylation occupancy spans four orders of magnitude, but the median ubiquitylation site occupancy is three orders of magnitude lower than that of phosphorylation. The occupancy, turnover rate, and the regulation of sites by proteasome inhibitors show strong interrelationships. These properties can discriminate signaling-relevant sites from the sites involved in proteasomal degradation. The sites strongly upregulated by proteasome inhibitors have a longer half-life, and the half-life increases with increasing protein length. Importantly, a previously unknown surveillance mechanism rapidly deubiquitylates all ubiquitin-specific E1 and E2 enzymes and protects them against bystander ubiquitylation accumulation. This work reveals general principles of ubiquitylation-dependent governance and offers conceptual insights into the dynamic regulation of the cell.

**Highlights:**

- Ubiquitylation site occupancy is 3 orders of magnitude lower than phosphorylation
- The highest 80% and the lowest 20% occupancy sites have distinct properties
- High occupancy sites are concentrated in the cytoplasmic domains of SLC proteins
- A dedicated mechanism prevevents ubiquitylation accumulation in E1s and E2s

## Introduction

To gain a quantitative understanding of dynamic operating systems, whether natural or man-made, two crucial requirements must be fulfilled. First, we need to identify and quantify the constituents of the system. Second, we must comprehend the mechanisms that dynamically control these constituents. A prime example of such a complex and dynamically regulated system is the mammalian cell, where proteins serve as the primary structural and functional building blocks. However, the function of proteins is reversibly regulated by posttranslational modifications (PTMs), which act as key switches in dynamical cellular control.

We have made significant progress in understanding the first requirement: we know the proteome-scale turnover rates and absolute abundance of mRNAs and proteins in diverse systems (Ghaemmaghami et al., 2003) (Buccitelli and Selbach, 2020; Cambridge et al., 2011; Herzog et al., 2017; Marguerat et al., 2012; Schwanhäusser et al., 2011; Taniguchi et al., 2010; Vogel and Marcotte, 2012; Zecha et al., 2018b). However, the second requirement remains largely unfulfilled: our knowledge of the absolute abundance and turnover rates of PTM sites, known as site occupancy or site stoichiometry, remain very limited. Indeed, site-specific turnover rates have not been quantified for any of the major eukaryotic PTMs on a global scale. PTMs impact proteins in a site-specific manner, therefore, without a quantitative understanding of individual regulatory sites, we cannot fully comprehend the dynamic regulation of a cell. After all, PTM sites are the main regulatory switches that dynamically regulate proteins and the cell.

Among the various PTMs, ubiquitination is the most complex regulatory mechanism that is specific to eukaryotes. In humans, ubiquitylation is governed by approximately 640 ubiquitylating enzymes and around 90 deubiquitylases (DUBs) that remove ubiquitin (Clague et al., 2015a; Komander et al., 2009). Ubiquitylation targets over 100,000 sites within the cell (Hansen et al., 2021; Kim et al., 2011; Steger et al., 2021; Wagner et al., 2011) Phosphorylation is the only other PTM that exhibits such broad scope and regulatory complexity, targeting over 100,000 sites and controlled by ∼540 kinases and ∼190 phosphatases (Brognard and Hunter, 2011; Chen et al., 2017; Wilson et al., 2018).

Although ubiquitylation has been extensively studied, the current research has primarily focused on measuring relative changes rather than absolute quantification. Site-specific occupancy has been measured for only a few sites (Kaiser et al., 2011; Li et al., 2019; Ordureau et al., 2018). These analyses offer valuable insights into specific sites analyzed but insufficient to provide a systems-scale perspective. Consequently, fundamental questions remain unanswered. For instance, what is the global occupancy of ubiquitylation sites? How quickly is ubiquitylation turned over at individual sites? Which sites have the highest occupancy? Which sites undergo the most rapid turnover? What is the half-life of different polyubiquitin linkages? How does proteasome activity affect ubiquitylation site turnover? Do sites involved in proteasomal and non-proteasomal functions differ in their abundance and/or turnover? Addressing these outstanding questions is crucial for understanding the basic principles underlying ubiquitylation-dependent regulation.

In this study, we conducted a comprehensive analysis of ubiquitylation site occupancy and half-life on a proteome-wide scale in human cells. By examining the occupancy of key PTMs, and comparing the turnover rates of ubiquitylation, mRNAs, and proteins, we discovered significant differences in the design principles of various regulatory layers. The work offers major new insights into the global attributes and principles of ubiquitylation-dependent cellular governance.

## Results

### Strategy for UB site occupancy measurement

Trypsin proteolysis of proteins modified by ubiquitin, NEDD8, or ISG15, leaves a Gly-Gly (GG) remnant attached to the modified lysine. This GG remnant is widely employed in mass spectrometry (MS) to map lysine residues harboring these PTMs. Although the GG remnant profiling does not distinguish between the three PTMs, most (>95%) GG modified sites are derived from ubiquitin-modified proteins (Kim et al., 2011). Therefore, for the sake of simplicity and consistency with existing literature, we refer to GG-modified sites as ubiquitylation sites in this study, as is commonly practiced.

To systematically quantify the occupancy of GG-modified sites, we employed a combination of the partial chemical modification (Prus et al., 2019; Weinert et al., 2015), Lys-ε-Gly-Gly (GG) remnant profiling (Kim et al., 2011; Wagner et al., 2011), and the serial dilution SILAC (SD-SILAC) methods (Weinert et al., 2015; Weinert et al., 2017). HeLa cells were chosen as the cell model for our analyses, given their wide usage in systems-scale studies. SILAC-heavy-labeled proteins were partially modified with GG using NHS-Gly-Gly-Boc (Xu et al., 2010) (**Figure S1**), and the degree of partial chemical GG modification (PC-GG) was measured using MS (**Figure S2**). A known amount of PC-GG-modified proteins (SILAC-heavy) was introduced into native proteins (SILAC-light). Following trypsin proteolysis, GG-modified peptides were enriched and quantified using MS. Site occupancy was calculated based on the relative abundance of native and chemically modified GG peptides (**Figure S1C**). To ensure accurate quantification of native peptides, we adjusted the amount of spiked-in proteins empirically, as an excessive or insufficient quantity can hinder the quantification process (Prus et al., 2019). In the initial experiment, PC-GG proteins were spiked in to achieve final occupancies of 3%, 1%, 0.1%, and 0.01%. Due to the quantification of fewer sites in the sample with 3% PC-GG spike-in, we adjusted the amount of spiked-in protein in all subsequent experiments to obtain final site occupancies of 1%, 0.1%, 0.01%, and 0.001%.

### Ubiquitylation occupancy is low but spans large dynamic range

We conducted a total of 7 biological replicates, where each replicate involved 4 serial dilutions of PC-GG-modified proteins. From these experiments, we identified a total of 34,247 sites and obtained SILAC ratios for 25,701 sites. When the abundance range of peptides is very large, quantification by MS becomes challenging. To ensure the accuracy of our quantification, we set a criterion that the measured SILAC ratios must agree (within two-fold variability) between at least two serial dilutions (Weinert et al., 2017). Using these accurately quantified sites, we determined the occupancy for 11,403 sites across 3,086 proteins (**Table S1**). The occupancy measurements exhibited strong correlation between replicates (r=0.82-0.97) (**Figure 1A**). Furthermore, 96.3% of the measurements showed less than a two-fold variation between replicates (median coefficient of variance 42.5%; median SD ± 0.0030%) (**Figure 1B**).

**Figure 1.**
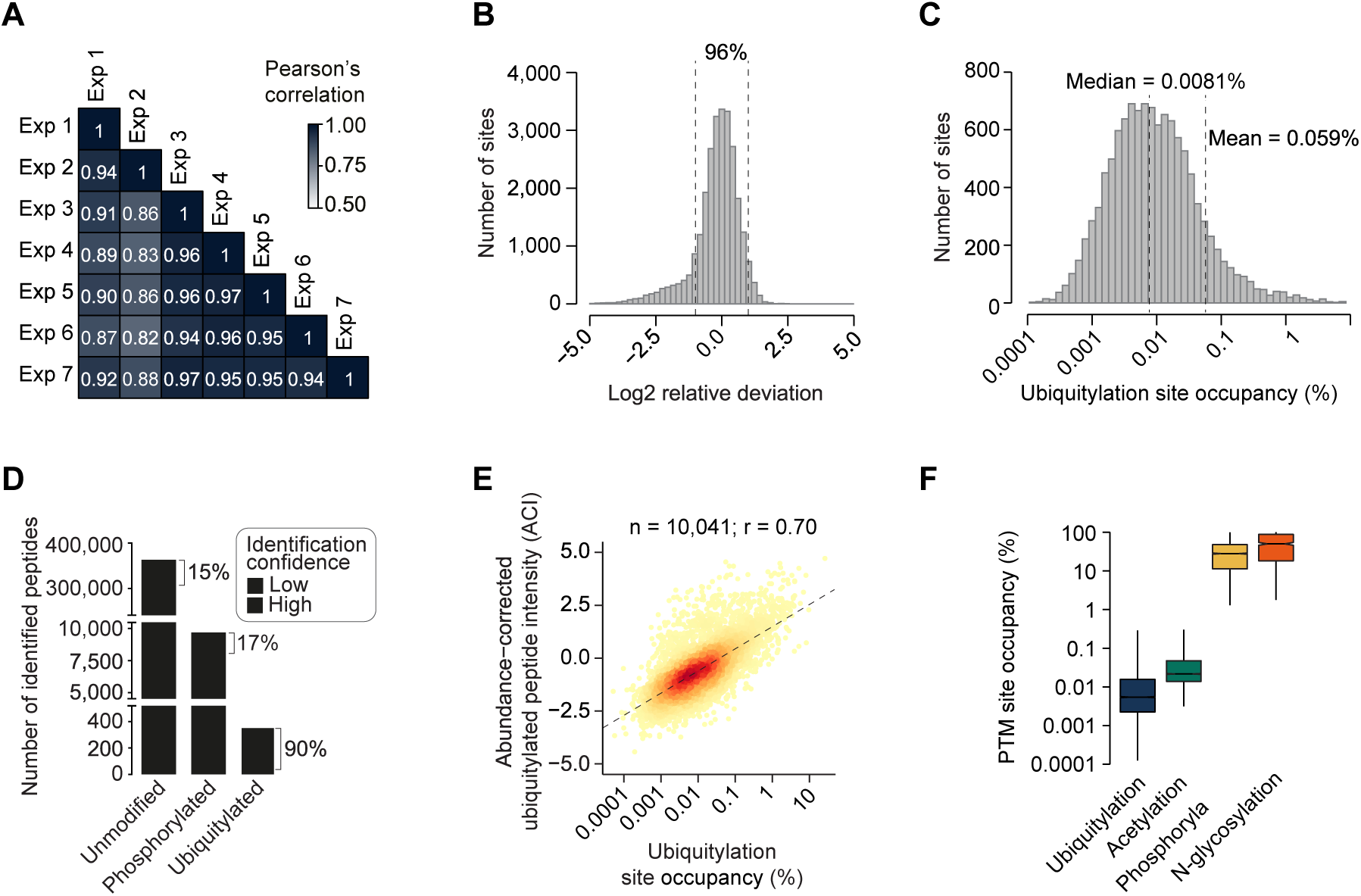
UB site occupancy is extremely low. (**A**) Reproducibility of UB site occupancy measurements. Shown is the pairwise correlation (Pearsońs) between seven different biological replicates. (**B**) The distribution of the relative deviation of site-specific ubiquitylation occupancy. Relative deviation is calculated as the standard deviation of ubiquitylation site occupancy in each experiment divided by the median occupancy from all experiments. (**C**) The distribution of the occupancies of ubiquitylation sites (n= 11,403) quantified in this study. Mean and median occupancy are indicated. (**C**) Ubiquitylated peptides are seldom detected in deep proteome analyses, without the affinity enrichment of ubiquitylated peptides. Shown is the number of unmodified, phosphorylated, and ubiquitylated peptides identified in a deep analysis of HeLa proteome (Bekker-Jensen et al., 2017), without the affinity enrichment for modified peptides. The number of peptides identified with standard database search (light grey bar; false discovery rate, 1%) and the number of high-confidence peptides retained after stringent filtering (dark black bar; see Methods). (**E**) Correlation between ubiquitylation site occupancy and protein abundance-corrected intensity (ACI) of ubiquitylated peptides. ACI is calculated by normalizing the intensity of GG-modified peptide with the abundance of the corresponding protein (see Methods). The number of analyzed ubiquitylation sites (n), Pearson’s correlation (r), and regression slope (dashed line) are shown. (**F**) The distribution of the site-specific occupancy of ubiquitylation (this study), acetylation (Hansen et al., 2019), phosphorylation (Olsen et al., 2010), and N-glycosylation (Yang et al., 2017). Occupancies of ubiquitylation, acetylation, and phosphorylation, were quantified in HeLa, while N-glycosylation occupancy was measured in HEK293. Box represents the interquartile range, the middle line denotes the median, and the whiskers indicate the minimum and maximum values excluding outliers.

Analyzing the data, we found that the median occupancy of GG-modified sites was 0.0081%, while the mean occupancy was 0.059% (**Figure 1C**). The skewed distribution of occupancy likely reflects the uneven distribution of protein abundance, where the median abundance is over ten times lower than the mean abundance (Bekker-Jensen et al., 2017). Only 1% (115/11,403) of the sites exhibited occupancy greater than 1%, and 1.9% of the sites had occupancy higher than 0.5%. These results indicate that the overall site occupancy of ubiquitylation, neddylation, and ISGylation is extremely low, despite the wide range of occupancy spanning over four orders of magnitude.

The low occupancy of GG-modified sites is corroborated by multiple lines of evidence. Firstly, when conducting a database search of the extensive HeLa proteome (Bekker-Jensen et al., 2017) without affinity enrichment of modified peptides, we found a large number of phosphorylated peptides but only a small number of GG-modified peptides. Moreover, the detected GG-modified peptides had a high rate of false positives (**Figure 1D**, **Supplementary Note 1**). After removing false positives, the remaining GG-modified sites in our dataset exhibited high occupancy and/or were found in highly abundant proteins (**Figure S3, A-B**).

Secondly, there is a strong correlation (r=0.70) between GG-modified site occupancy and the protein abundance corrected intensity of GG-modified peptides (ACI) (Weinert et al., 2015) (**Figure 1E, Supplementary Note 2**). This correlation is comparable to, or even higher than, the maximum empirical correlation (r=0.63-0.65) observed between unmodified peptide intensity and the abundance of the corresponding protein (**Figure S3C**). Since GG-modified site occupancy and ACI are determined independently of each other, the high correlation between them strongly supports the accuracy of our measurements.

Thirdly, empirical occupancy measurements align with theoretical estimates. Based on our calculations, we estimate that ubiquitin accounts for approximately 1.1% of total protein molecules in HeLa (**see Methods**). Considering the number of ubiquitin molecules in a HeLa cell, the total number of proteins, the count of ubiquitylated lysines, and the fraction of ubiquitin conjugated to proteins, we derive a median theoretical occupancy for ubiquitylation sites of 0.0085% (**Supplementary Note 3**). This estimation closely matches our experimentally measured occupancy (median, 0.0081%).

### Different PTMs have vastly different occupancies

Mammalian proteins are modified with diverse types of PTMs, raising the question of how the occupancy of ubiquitylation compares to other PTMs. To address this question, we compared the occupancy of four commonly observed PTMs: ubiquitylation, phosphorylation (Olsen et al., 2010), acetylation (Hansen et al., 2019), and N-glycosylation (Yang et al., 2017) (**Figure 1F**). Among these PTMs, ubiquitylation exhibits the lowest site occupancy. In fact, the median occupancy of ubiquitylation is more than three orders of magnitude lower than that of phosphorylation (28%). On the other hand, N-glycosylation demonstrates the highest occupancy, with many sites being modified almost to full occupancy (Yang et al., 2017; Zielinska et al., 2010). These very large differences in occupancy imply that distinct PTMs operate through distinct mechanistic principles.

### Ubiquitylation is turned over kinetically and site selectively

To reconcile the seemingly contradictory low occupancy with the vast regulatory role of ubiquitylation, we investigated the site-specific half-life of ubiquitylation. To the best of our knowledge, no previous study has quantified the in vivo half-life of a widely occurring eukaryotic PTM in a site-specific manner on a global scale. Although one study examined the acetylation site half-life using metabolic labeling, the accuracy of measurements was greatly compromised due to the long time periods (several hours) required for the incorporation of labeled metabolite precursors into PTMs (Kori et al., 2017) (**Supplementary Note 4**). In the ubiquitylation and neddylation systems, the E1 enzymes (UBA1 and UBA6 for ubiquitin; and NAE for NEDD8) function at the top, and inhibiting E1 enzymes completely abolish catalysis of these PTMs. To quantify site-specific deubiquitylation rates, we used TAK-243 (MLN7243), which inhibits both UBA1 and UBA6 (Hyer et al., 2018). We determined the concentration that fully inhibits ubiquitylation and utilized a saturating concentration (100µM) for all proteomic analyses (**Figure S4, A-B, Supplementary Note 5**). It’s worth noting that at 10 µM, TAK-243 also strongly inhibits neddylation (Hyer et al., 2018), thus in our analyses, we inhibited both ubiquitylation and neddylation. In time-course experiments, cells were treated with TAK-243 for 5, 10, 30, and 60 minutes, and the change in ubiquitylation was quantified using SILAC-based MS (**Figure S4C**). From 5 biological replicates, we cumulatively identified 24,112 sites. TAK-243 treatment led to a time-dependent decrease in global ubiquitylation (**Figure 2A, S4 B**). Sites quantified across all time points (13,833 sites occurring in 4,065 proteins) were used to calculate site-specific deubiquitylation rates (half-life, t_1/2_) (**Table S2**). The site-specific half-life was determined by applying either the exponential decay model (7,870 sites) or the exponential-plateau model (5,963 sites) to the kinetic profiles (**see Methods**). We classified ubiquitylation sites into four categories: ‘Very fast’, ‘Fast’, ‘Slow’, and ‘Very slow’, corresponding to half-lives of <5, 5-15, 15-60, and >60 minutes, respectively (**Figure 2B-D**). Over 22% of sites exhibited a half-life of less than 5 minutes, 45% had a half-life of less than 10 minutes, and 67% had a half-life of less than 30 minutes. Approximately 14% of sites showed no significant decrease in ubiquitylation within 1 hour of TAK-243 treatment, making it challenging to accurately calculate their half-life. For these sites, we indicated their half-life as >120 minutes (twice the length of the longest treatment time point in our analyses). The rapid half-life of ubiquitylation contrasts sharply with the half-life of proteins, where 99% of proteins have a half-life longer than 30 minutes (Li et al., 2021).

**Figure 2.**
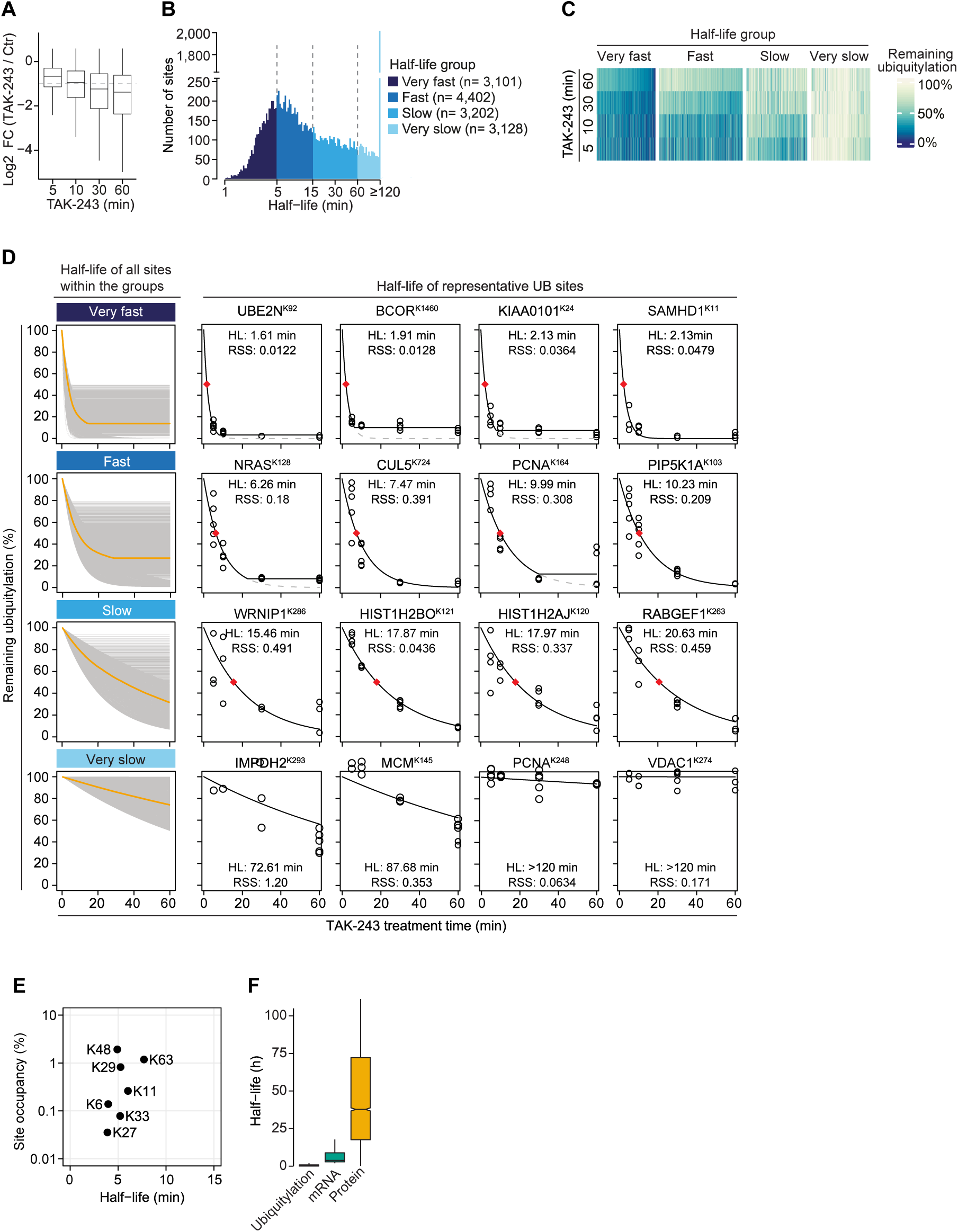
Ubiquitylation site half-life is very short. (**A**) E1 inhibition causes a time-dependent decrease in global ubiquitylation. Shown is the change in ubiquitylation after E1 inhibition by TAK-243 (100µM) for the indicated time points. Ubiquitylation was quantified by mass spectrometry using the GG remnant profiling approach. Box represents the interquartile range, the middle line denotes the median, and the whiskers indicate the minimum and maximum values excluding outliers. (**B**) The distribution of ubiquitylation site half-life. Ubiquitylation sites are grouped into the indicated four categories (Very fast, Fast, Slow, and Very slow), and the number of sites (n) in each group is indicated. (**C**) The heatmaps show temporal change in site-specific ubiquitylation after TAK-243 treatment for the indicated time. (**D**) Shown are the deubiquitylation kinetics of UB sites in the indicated half-life groups. The left panels show combined profiles of all UB sites in each of the indicated groups, and the orange lines indicate the general trend within each group. From each half-life group, kinetic profiles of four representative UB sites are shown (right panels). The sites for which the reduction in ubiquitylation reaches a plateau, the dashed lines indicate an extrapolation of the exponential part of the function. The indicated half-life (t_1/2_, indicated by the red diamond symbol) values were calculated based on the exponential part of the decay curve. The root sum squares (RSS) indicate deviation between experimentally measured values and values expected from the fitted decay. (**E**) Shown are the half-life and occupancy of the indicated polyubiquitin linkage types. (**F**) Ubiquitylation is turned over much shorter than mRNA and protein. Shown is the distribution of the half-lives of ubiquitylation, mRNA (Tani et al., 2012), and protein (Zecha et al., 2018) in HeLa.

The global half-life of ubiquitylation sites is remarkably short, with a median of 12 minutes. However, the half-lives of individual sites exhibit significant variation spanning over two orders of magnitude, ranging from less than 1 minute to more than 120 minutes. Site-specific ubiquitylation serves diverse signaling functions and undergoes dynamic regulation in response to stimuli. For instance, histone H2A^K119^ and H2B^K120^ ubiquitylation respectively regulate transcription repression and activation (Weake and Workman, 2008). Ubiquitylation of PCNA^K164^ controls translesion synthesis, while PCNA-associated factor PAF15^K24^ (KIAA0101^K24^) ubiquitylation promotes replication-associated DNA methylation (Hoege et al., 2002; Nishiyama et al., 2020; Povlsen et al., 2012). Other examples include CUL5^K724^ neddylation required for the activation of cullin-RING family ubiquitin ligases (Duda et al., 2008), NRAS^K128^ induction upon B cell receptor activation (Satpathy et al., 2015), PIP5K1A^K103^ dynamic regulation upon Salmonella infection (Fiskin et al., 2016), and VDAC1^K274^ increase during mitophagy, implicated in inhibiting apoptosis (Ham et al., 2020; Ordureau et al., 2018).

To our knowledge, the in-vivo turnover rates of these specific sites have not been quantified. Our findings reveal that the half-lives of these sites vary greatly, ranging from <2 minutes to >120 minutes (**Figure 2D**). It’s worth noting that the rapid turnover of known ubiquitylation sites, as well as the neddylation site CUL5^K724^, confirms the effective inhibition of both ubiquitylation and neddylation in our assays. Moreover, within the same proteins, the half-lives vary site-specifically. For example, the half-lives of PCNA^K164^ versus PCNA^K248^, NRAS^K128^ versus NRAS^K5^, and SAMHD1^K11^ versus SAMHD1^K446^ differ ∼10-50 times (**Figure S4D**). Notably, the fastest turnover sites within these proteins exhibit the highest occupancy, highlighting the remarkable selectivity of ubiquitin ligases and DUBs. Interestingly, all polyubiquitin linkages undergo rapid turnover (half-life ∼4-8 minutes) (**Figure 2E**), indicating that fast turnover is not limited to a specific type of ubiquitin linkage. The similar turnover rates of ubiquitin linkages suggest that differences in the abundance of polyubiquitin linkages primarily result from variations in catalytic efficiency and/or the number of E3 ligases catalyzing different linkages, rather than differences in their deubiquitylation rates. In conclusion, these results demonstrate that while the global half-life of ubiquitylation is short, the half-life of individual sites displays considerable variability, suggesting large differences in the kinetics of ubiquitylation-regulated processes.

### Ubiquitylation is turned over much faster than mRNA and protein

Eukaryotic systems are regulated through various levels, including transcriptional, translational, and posttranslational mechanisms. To gain insights into the comparative dynamics of these regulatory layers, we compared the half-lives of ubiquitylation with the half-lives of mRNA and protein (Cambridge et al., 2011; Doherty et al., 2009; Herzog et al., 2017; Pratt et al., 2002; Schwanhüusser et al., 2011; Tani et al., 2012; Yang et al., 2003; Zecha et al., 2018b). In comparison to ubiquitylation, the half-life of mRNAs is much longer, with a median range of approximately 4 to 9 hours. The half-life of proteins is even longer, with a median range of approximately 37 to 46 hours, making it approximately 200 times longer than the half-life of ubiquitylation (**Figure 2F**). These substantial disparities in the turnover rates of mRNA, protein, and ubiquitylation indicate that these distinct regulatory layers are tailored to provide regulation at different timescales.

### Global deubiquitylation continues without proteasome activity

The proteasome plays a crucial role as a central hub for ubiquitin recycling, and the deubiquitylation of proteasome substrates is coupled to proteasome activity (Verma et al., 2002; Yao and Cohen, 2002). To investigate the importance of proteasome activity in global deubiquitylation, we examined deubiquitylation in cells treated with only the E1 inhibitor or in combination with the proteasome inhibitor MG-132. We observed rapid deubiquitylation of proteins even when the proteasome was inhibited (**Figure 3, A-B**, **Table S3**). This finding indicated a decoupling between global deubiquitylation and proteasome activity. To further confirm this decoupling, we pre-treated cells with two distinct proteasome inhibitors (MG-132 and bortezomib (BTZ)) to enhance the ubiquitylation of proteasome substrates. We then inhibited E1 to monitor the kinetics of deubiquitylation (**Figure S3C**). As expected, MG-132 and BTZ effectively increased ubiquitylation levels (**Figure 3D, Table S3**). However, the subsequent addition of TAK-243 drastically reduced MG-132 and BTZ-induced ubiquitylation (**Figure 3E**). This finding suggests that when proteasome activity is compromised, the deubiquitylation of proteins destined for proteasomal degradation can occur independently of proteasome function. Conceivably, this decoupling phenomenon could potentially help alleviate proteostasis stress by providing a backup mechanism for ubiquitin recycling, thereby supporting non-proteasomal ubiquitylation functions.

**Figure 3.**
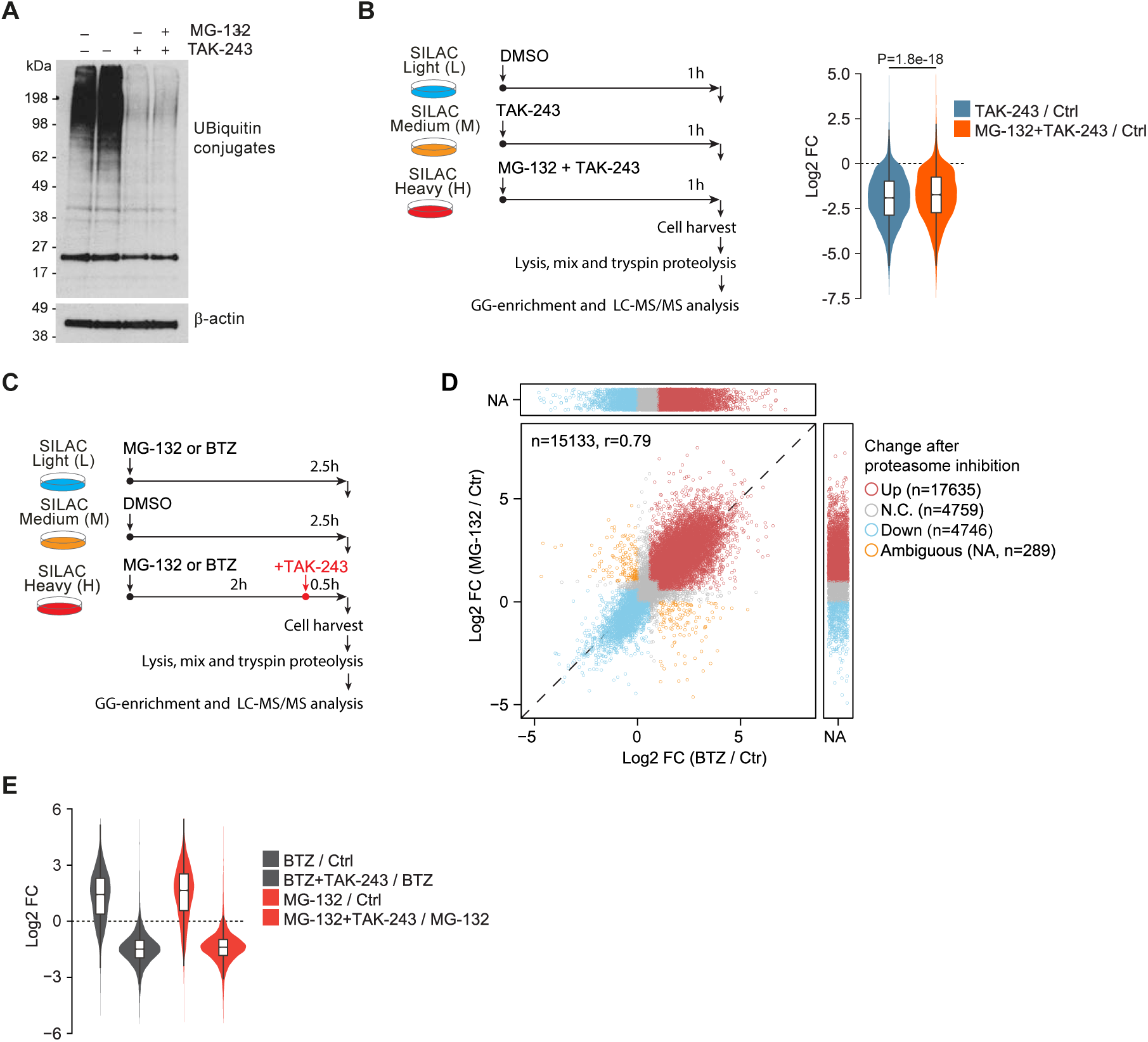
Global protein deubiquitylation can be decoupled from the proteasome activity. (**A**) The relative levels of ubiquitin conjugates in cells treated without or with the indicated combinations of TAK-243 (100µM) and MG-132 (200µM). Beta-actin was used as a loading control (n=2). (**B**) Schematic (left panel) of ubiquitylation quantification after inhibition of E1, with or without MG-132. SILAC-labeled cells were treated (1h) with solvent (DMSO), or E1 inhibitor (100µM TAK-243), or the combination of E1 inhibitor and a proteasome inhibitor (100µM TAK-243 + 200µM MG-132). Changes in ubiquitylation were quantified through the GG remnant profiling. Ubiquitylation is strongly decreased after the inhibition of E1 alone, and the simultaneous inhibition of E1 and the proteasome (right panel). The experiment was performed in 3 biological replicates. P-values were calculated using a two-sided Wilcoxon rank-sum test. (**C**) Schematic of SILAC-based quantification of deubiquitylation of proteasome inhibitor regulated sites. SILAC-labeled cells were treated (2.5h) with solvent (DMSO), or proteasome inhibitors MG-132 (200µM) or BTZ (100µM). For sequential treatment of cells with proteasome inhibitor and E1 inhibitor, the cells were pre-treated (2h) with MG-132 (200µM) or BTZ (100µM), and the E1 inhibitor (100µM TAK-243) was added 30 minutes prior to harvest. Changes in ubiquitylation were quantified through the GG remnant profiling by mass spectrometry. (**D**) Proteasome inhibition by MG-132 and BTZ causes widespread changes in ubiquitylation. Change in ubiquitylation were quantified as described in the panel C, and the correlation (Pearsońs) between MG-132- and BTZ-induced ubiquitylation change is shown. Based on the change in ubiquitylation, sites were classified as upregulated (Up), downregulated (Down), not changing (N.C.), or ambiguous (NA) (see Methods for further details on the classification of sites). Sites classified as ambiguous (NA) were not regulated consistently between MG-132 and BTZ and were not used for further analyses. Correlation (Pearsonśs) and the number of Up, Down, N.C., and NA sites are indicated. (**E**) Proteasome inhibition-induced ubiquitylation is decreased following E1 inhibition. Change in ubiquitylation were quantified as described in the panel C. The relative changes in ubiquitylation changes were quantified after treating the cells with one of the indicated proteasome inhibitors, or after sequential treatment of cells with the indicated proteasome inhibitors and TAK-243, as depicted in the schematic (panel C). The violin plots show the SILAC ratio of UB sites after the indicated treatments.

### High occupancy sites are concentrated in plasma membrane receptors

Next, we asked which proteins harbor the highest occupancy UB sites. Among the top sites with the highest occupancy (>1%), we identified well-known ubiquitylation and neddylation sites of functional importance. These included histone H2A^K119^, UB^K48^, UB^K63^, CUL5^K724^, RHOA^K135^, and CXCR4^K327,K331^ (**Table S1**). For instance, H2A^K119^ ubiquitylation is associated with transcription repression (Weake and Workman, 2008), UB^K48^ and UB^K63^are involved in diverse processes, CUL5^K724^ neddylation regulates cullin-RING ligases (Duda et al., 2008), and CXCR4^K327,K331^ controls the lysosomal sorting of the receptor (Marchese and Benovic, 2001).

Notably, sites with high occupancy (>0.5%) were significantly enriched in Gene Ontology (GO) terms related to the plasma membrane (PM) and transmembrane (TM) transporter activity (**Figure 4A, Table S4**). In fact, more than half of the top 100 sites with the highest occupancy were found in TM receptors associated with the plasma membrane (**Figure 4B**). Overall, TM proteins exhibited higher occupancy compared to non-TM proteins. Within TM proteins, the solute carrier family (SLC) proteins showed higher occupancy compared to non-SLC proteins (**Figure 4D**). The median site occupancy of PM-associated SLCs (0.96%) was more than 100 times greater than the median occupancy of all sites combined. While PM-associated non-SLCs also displayed higher-than-average occupancy, their occupancy was much lower than that of sites in PM-bound SLCs (**Figure 4D**). SLCs encompass a diverse group of transporters localized in various subcellular compartments (Pizzagalli et al., 2021). High occupancy sites were specifically enriched among PM-localized SLCs (**Figure 4D**). Among the top 100 sites with the highest occupancy, 44% were found on PM-bound SLCs, despite these proteins constituting only 0.85% of the total sites in the occupancy dataset (**Figure 4C**). This observation highlights that high occupancy is a characteristic feature of PM-associated SLCs as a specific group.

**Figure 4.**
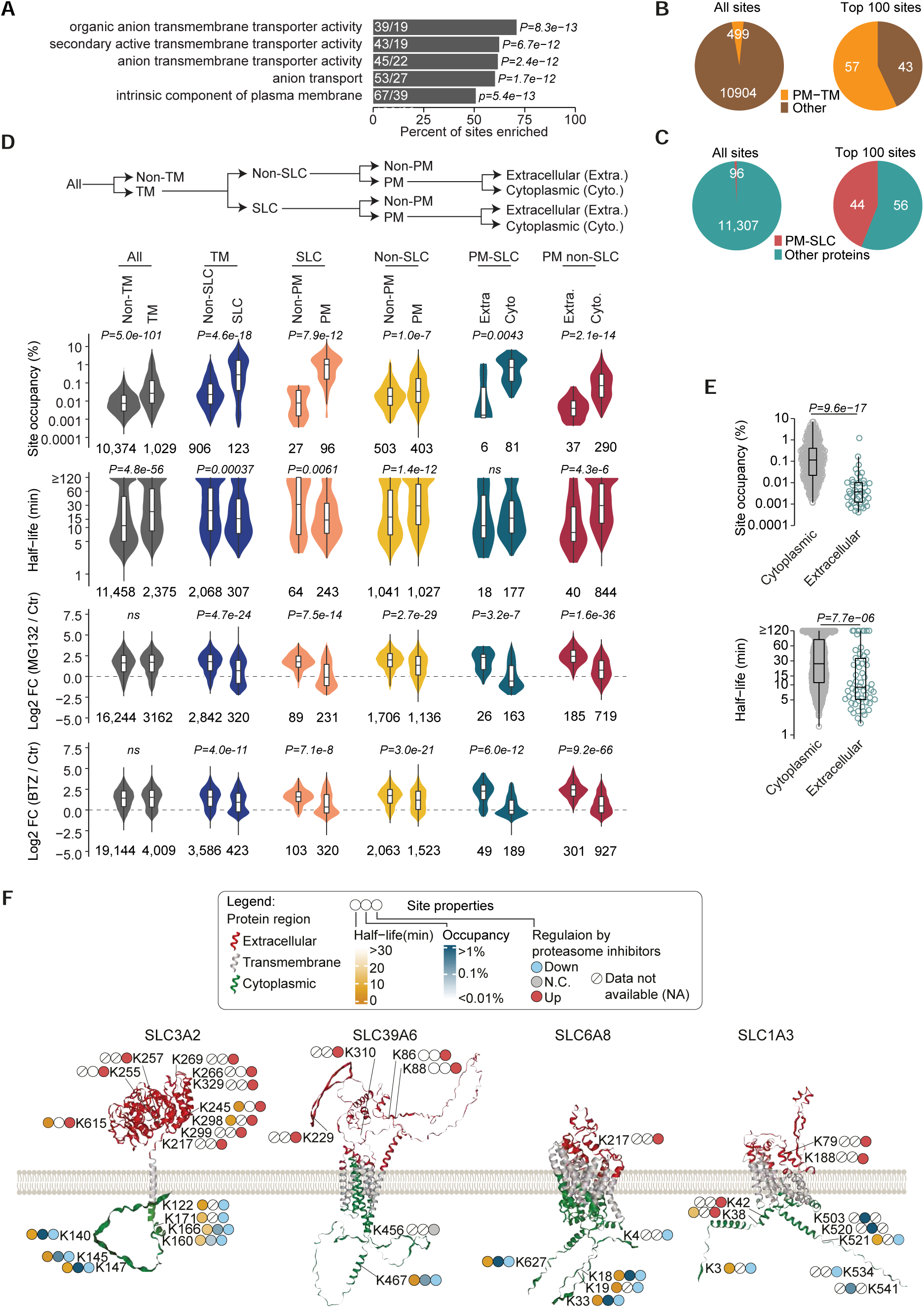
High occupancy ubiquitylation sites are concentrated in the intracellular domains of the plasma membrane receptors. (**A**) Gene Ontology (GO) enrichment analysis of proteins harboring high occupancy (>0.5%, n=218) sites. Shown are the selected GO terms that were most significantly enriched. Percent site enrichment indicates the fraction of high occupancy sites in proteins enriched in each GO term. Within each GO category, the number of high occupancy sites and the number of ubiquitylated proteins are shown (n, UB sites/ n, proteins). P-values indicate the Bonferroni-corrected statistical significance. (**B-C**) Shown is the fraction of UB sites identified in the plasma membrane transmembrane (PM-TM) proteins (**B**) and plasma membrane SLC (PM-SLC) proteins (**C**), among all sites (left panel), and the 100 highest occupancy sites (right panel). (**D**) Distinct properties of UB sites in transmembrane receptors. Proteins were grouped into non-transmembrane and transmembrane categories, and within the transmembrane category, the proteins were grouped further groped into the indicated categories, as indicated (top panel). Within each category, the distribution of UB site occupancy, half-life, and regulation upon proteasome inhibition (MG-132 or BTZ, 2.5h) was analyzed (bottom panels). Box represents the interquartile range, the middle line denotes the median, and the whiskers indicate the minimum and maximum values excluding outliers. The number of UB sites (n) in each group and P-values for comparison between the groups are indicated. P-values were calculated using a two-sided Wilcoxon rank-sum test (ns, P >0.05). TM: transmembrane, PM: plasma membrane, SLC: solute carrier family proteins, Extra.: extracellular protein domain, Cyto.: cytoplasmic protein domain. (**E**) Shown is the distribution of occupancy (left panel) and the half-life (right panel) of the UB sites occurring in the cytoplasmic and extracellular regions of plasma membrane-associated transmembrane proteins. P-values were calculated using a two-sided Wilcoxon rank-sum test. (**F**) The sites in the intracellular (cytoplasmic) and extracellular domains of the plasma membrane SLCs show different properties. Shown are the AlphaFold-predicted structures of the indicated SLC proteins, as well as the properties of the identified ubiquitylation sites. Different topological domains (cytoplasmic, extracellular, transmembrane) are color-coded as indicated. The identified ubiquitylation sites, their occupancy, half-life, and regulation by proteasome inhibition are shown.

### Extracellular and cytoplasmic sites in PM receptors have distinct attributes

The remarkably high occupancy observed in PM-associated receptors prompted us to delve deeper into their characteristics. We observed a notable difference when comparing non-PM SLCs and PM-SLCs: ubiquitylation levels increased globally after proteasome inhibition in non-PM SLCs, whereas this global increase was not observed in PM-SLCs (**Figure 4D**). Upon closer examination of various PM-SLCs, we discovered distinct properties between sites located in the cytoplasmic and extracellular regions. Specifically, sites in the cytoplasmic domains exhibited higher occupancy and were unaffected by proteasome inhibitors, while sites in the extracellular regions showed a significant increase in response to proteasome inhibitors. Additionally, these extracellular sites generally displayed lower occupancy or were undetectable in the occupancy measurements (**Figure 4F**). Similar patterns were observed in several PM-associated non-SLC receptors (**Figure S5**).

On a global scale, when considering PM-associated TM proteins, we found that sites in the ctyplasmic regions had higher occupancy and longer half-lives compared to sites in the extracellular regions (**Figure 4E**). This suggests that ubiquitylation in the extracellular and intracellular domains occurs in different protein pools and serves distinct biological functions. PM-associated TM proteins are synthesized in the endoplasmic reticulum (ER) and subsequently transported to the membrane. The ubiquitylation of sites in the extracellular domains likely involves the ER-associated degradation (ERAD) machinery, which functions in quality control by targeting improperly folded receptors for proteasomal degradation. On the other hand, sites in the intracellular domains likely occur in mature proteins and serve non-proteasomal functions, such as the regulation of receptor trafficking.

### Ribosome-associated quality control sites have distinctively high occupancy

Non-proteasomal ubiquitylation plays a critical role in ribosome-associated protein quality control (RQC), which is activated by ribosome stalling and collisions between ribosomes (Dougherty et al., 2020; Filbeck et al., 2022). This process triggers site-specific ubiquitylation at specific sites on the 40S ribosomal subunit, including RPS10^K107, K138, K139^, RPS20^K4,K8^, RPS3^K214^, RPS7^K85^, and RPS27A^K107, K113^ (hereafter we collectively refer to these sites as RQC-associated sites) (Garzia et al., 2017; Ikeuchi et al., 2019; Montellese et al., 2020 ; Oltion et al., 2022). However, the occupancy and half-life of these RQC-associated sites have not been previously quantified. We aimed to determine whether the properties of these signaling-relevant sites could distinguish them from other ribosomal sites.

We identified a total of 863 sites in ribosomal proteins and quantified the occupancy and half-life for the majority of these sites (**Figure 5A**). Comparing ribosomal sites to non-ribosomal sites, we observed that ribosomal sites have significantly shorter half-lives, lower occupancy, and are less upregulated by proteasome inhibitors (**Figure 5B**). Occupancy was quantified for over 700 ribosomal sites, including all of the RQC-associated sites. Strikingly, all RQC-associated sites exhibited distinctively high occupancy. Among the 9 RQC-associated sites, 8 ranked among the top, with only RPS10^K139^ ranking slightly lower at position #17 (**Figure 5C**). Additionally, we detected moderately high occupancy for three other sites in the 40S subunit (RPS17^K107^, RPS20^K30^, and RPS12^K129^). These sites, like the other RQC-ubiquitylated subunits, localize near the interface of collided ribosomes (**Figure 5F**), and their ubiquitylation increases after ribosome stalling (52), indicating that they may also be ubiquitylated by RQC-associated ligases, albeit less efficiently. Importantly, RQC-associated sites exhibit short half-lives and their ubiquitylation is not increased by proteasome inhibition (**Figure 5 D-E**), supporting their non-proteasomal regulatory role. The short half-life of these key RQC-associated sites suggests that their high occupancy is not solely due to slower turnover.

**Figure 5.**
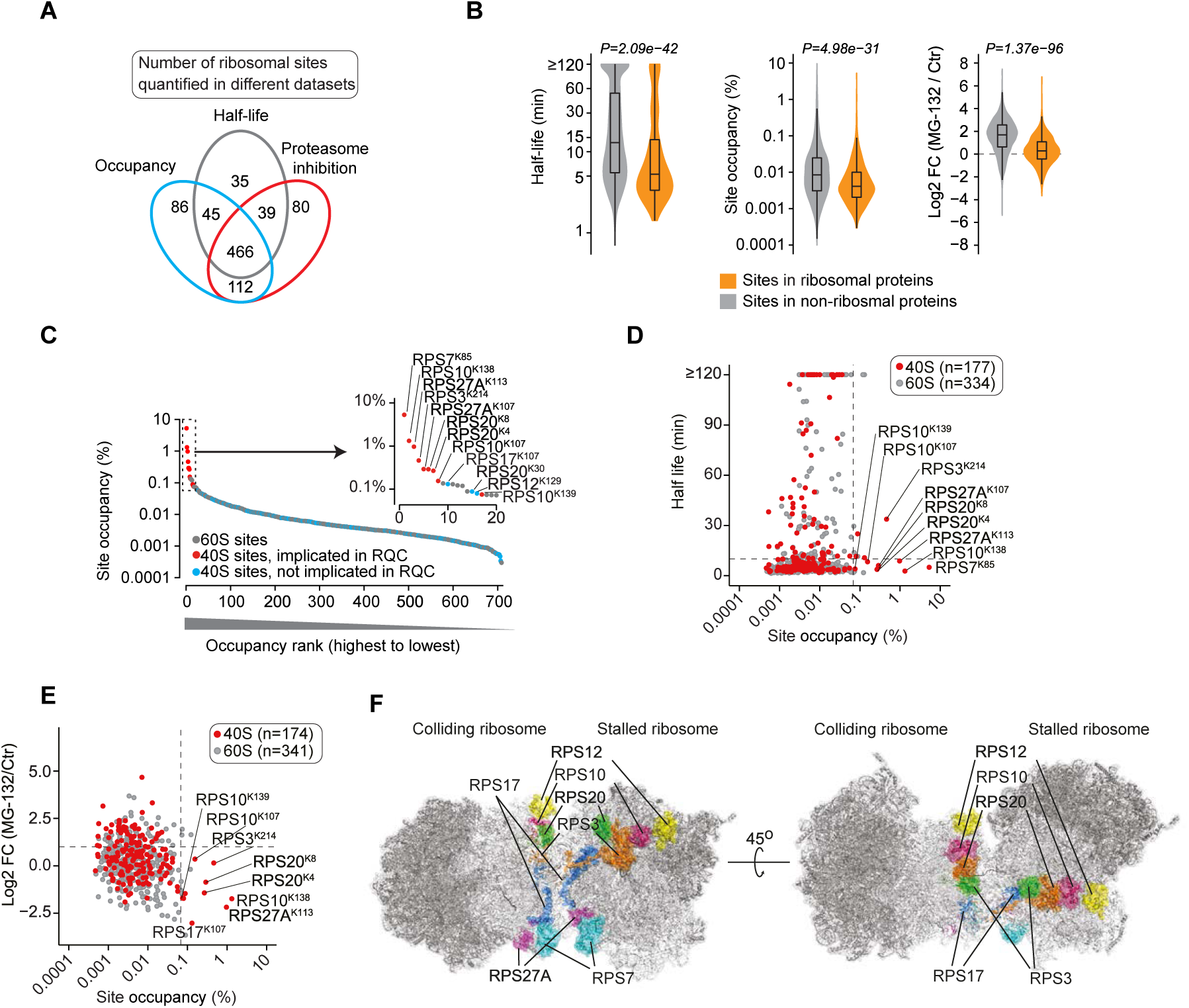
Sites involved in ribosome-associated quality control (RQC) show high occupancy and rapid turnover. (**A**) The overlap between the number of UB sites in ribosomal proteins quantified in the half-life, occupancy, and proteasome inhibition datasets. (**B**) Shown is the distribution of site half-life, occupancy, and their regulation by MG-132 (2.5h) in the ribosomal and non-ribosomal proteins. Box represents the interquartile range, the middle line denotes the median, and the whiskers indicate the minimum and maximum values excluding outliers. P-values were calculated using a two-sided Wilcoxon rank-sum test. (**C**) A distinctively high occupancy of the sites involved in ribosome-associated quality control (RQC). Shown is the rank plot of occupancies of all ubiquitylation sites mapped in ribosomal proteins. Sites are color coded as indicated, and the sites previously implicated in RQC (Garzia et al., 2017; Ikeuchi et al., 2019; Montellese et al., 2020 ; Oltion et al., 2022) are denoted. (**D-E**) Shown is the relationship between ribosomal UB site occupancy and half-life (**D**) and between occupancy and regulation by MG-132 (2.5h) (**E**). Sites in the 40S and 60S subunits are color coded as indicated and known RQC-associated sites are denoted. (**F**) Shown are the locations of the 40S subunits that harbor the top one dozen highest occupancy (>0.075%) sites in the disome structure. The structure of the budding yeast disome was obtained from RCSB Protein Data Bank (PDB ID: 6I7O) (*54*).

In addition to RQC-associated sites, distinct regulation patterns were observed among ribosomal subunits (**Figure S6**). For instance, RPL10 is the only subunit in which all sites display a short half-life and their ubiquitylation increases upon proteasome inhibition. RPL10 is one of the last subunits to assemble into the 60S subunit (Kruiswijk et al., 1978). The properties of RPL10 UB sites support the idea that late-assembling subunits of multi-protein complexes are targeted for non-exponential degradation by the proteasome until they are fully incorporated into mature complexes (McShane et al., 2016; Yanagitani et al., 2017). Taken together, these findings highlight that ribosome ubiquitylation is regulated in a site- and protein-specific manner, and the properties of these sites can effectively distinguish functionally relevant sites within this large and complex structure.

### E1 and E2 enzymes are deubiquitylated ultra-rapidly and site-indiscriminately

Which proteins are deubiquitylated slowly and which proteins are deubiquitylated rapidly? We found that sites with long half-lives were enriched in proteins associated with the Gene Ontology (GO) terms ‘ATP-dependent activity’, ‘ATP-dependent activity, acting on DNA’, and ‘vesicle lumen’ (**Figure 6A**). On the other hand, sites with very fast half-lives were highly enriched for the GO term ‘ubiquitin-conjugating enzyme activity’ (**Figure 6A**).

**Figure 6.**
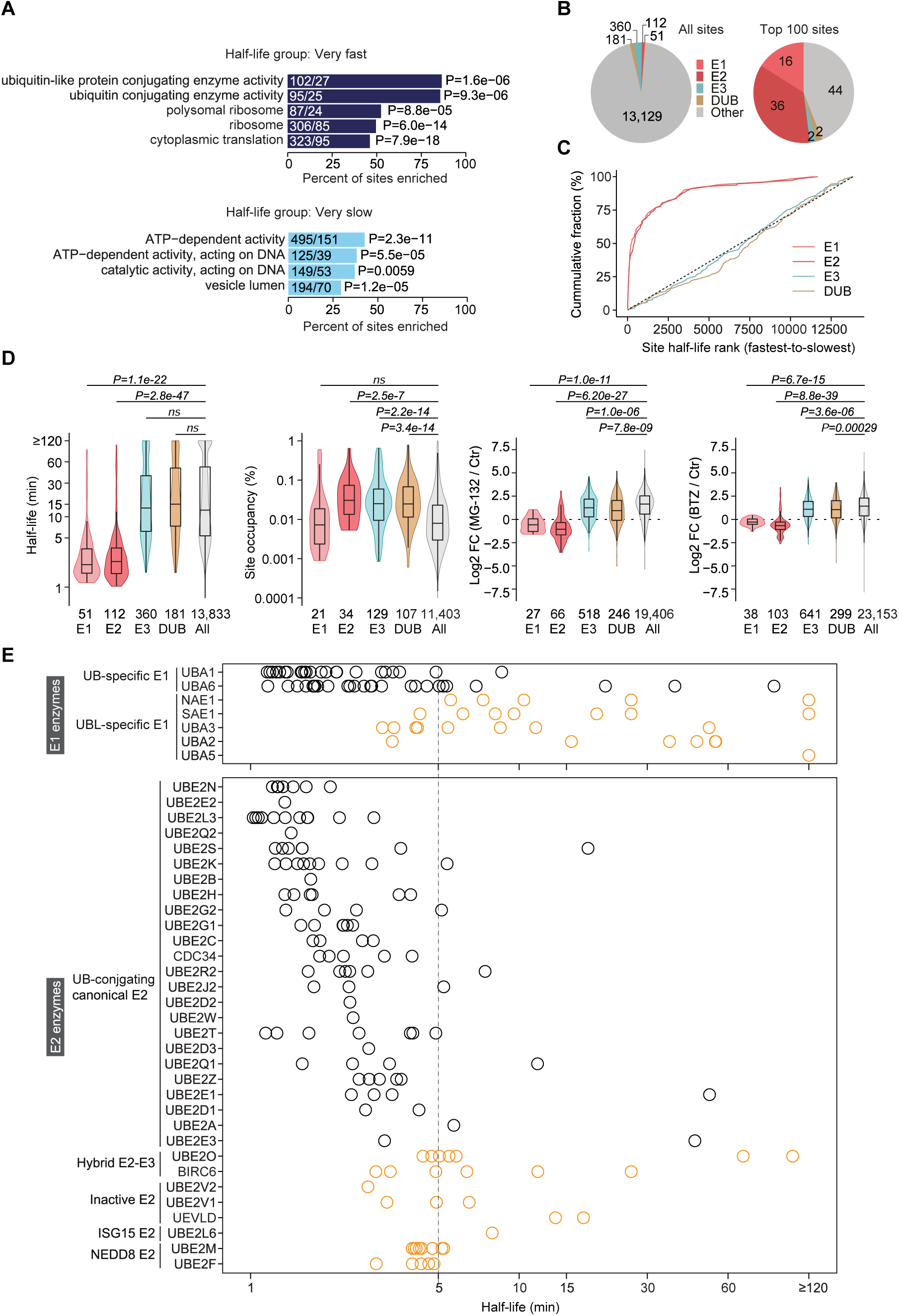
The ubiquitin-specific E1 and E2 enzymes are deubiquitylated extremely rapidly. (**A**) Shown are the Gene Ontology (GO) terms enriched in the proteins harboring ubiquitylation sites with very rapid (≤5 min; top panel) and very slow (≥60 min; bottom panel) half-lives. Within each GO category, the number of significantly enriched sites and the number of ubiquitylated proteins are shown (n, UB sites/ n, proteins). Percent site enrichment indicates the fraction of very rapid half-life sites in proteins assigned to a given GO term. P-values indicate the Bonferroni-corrected statistical significance. (**B**) Shown is the fraction of UB sites mapping to the indicated groups of proteins, among all quantified sites (left panel), and the top 100 sites with the fastest half-life (right panel). (**C**) UB sites are rank-ordered based on the half-life, and the cumulative fraction of UB sites occurring in the indicated group of proteins is shown. (**D**) Ubiquitin-specific E1-E2s, but not E3s and deubiquitylases (DUBs), are rapidly deubiquitylated. Shown is the distribution of UB site half-life, occupancy, and regulation by MG-132 and BTZ, in the indicated groups of proteins. Box represents the interquartile range, the middle line denotes the median, and the whiskers indicate the minimum and maximum values excluding outliers. The number of sites (n) in each group and P-values for comparison between groups are indicated. P-values were calculated using a two-sided Wilcoxon rank-sum test (*ns*, P>0.05). (**E**) Shown is the half-life of UB sites in the indicated E1 (upper panel) and E2 (lower panel) enzymes. Circles denote individual UB sites in the given protein. E1s and E2s are classified as specified.

Interestingly, only the ubiquitin-specific E1s and E2 enzymes exhibited rapid half-lives, while sites in E3s and deubiquitylating enzymes (DUBs) had average half-lives (**Figure 4 C-D**, **Figure S11B**). It is noteworthy that the occupancy of E2 sites did not significantly differ from that of sites in E3s and DUBs, and the sites in E1-E2s were not upregulated by proteasome inhibitors (**Figure 4D**).

Specifically, in ubiquitin-specific E1-E2s, 85% (138/163) of sites had a half-life of less than 5 minutes, with a median half-life of just 2.3 minutes. Although sites in E1-E2s constituted only 1.2% of the total quantified sites, they accounted for more than half of the top 100 sites with the shortest half-lives (**Figure 4B**). Moreover, within the E1 class, sites in the ubiquitin-activating E1s (UBA1 and UBA6) exhibited much faster half-lives compared to sites in E1s involved in the conjugation of ubiquitin-like modifiers (UBLs) (**Figure 6E**). These findings collectively identify ubiquitin-specific E1-E2s as the most rapidly (de)ubiquitylated protein class and suggest the existence of a dedicated mechanism that constantly and indiscriminately deubiquitylates E1-E2s at virtually all sites.

### Ubiquitylation occupancy, half-life, and proteasome regulation are interrelated

To gain a comprehensive understanding, we integrated datasets of UB site occupancy, half-life, and regulation by proteasome inhibition (**Table S5**), and examined the interrelationships these properties. Out of the sites quantified after proteasome inhibition (n=27,137), we were able to determine occupancy for 7,433 sites (**Figure S7A**). Interestingly, sites that were upregulated by MG-132 exhibited longer half-lives but lower occupancy (**Figure S7B**). Additionally, as the degree of upregulation by MG-132 increased, the proportion of sites quantified in terms of half-life and occupancy decreased linearly (**Figure S7C**). This suggests that sites showing a substantial relative increase after proteasome inhibition have very low occupancy in unperturbed cells, and remain below the detection limit of MS.

To explore the relationship between occupancy and half-life, we ranked and grouped (n=200) sites based on their occupancy. Within each group, we analyzed the fractions of sites belonging to different half-life categories. Remarkably, with increasing occupancy, the fraction of sites with a ‘Very fast’ half-life decreases almost linearly, while the proportion of sites with a ‘Very slow’ half-life increases (**Figure 7A**). However, this trend levels off for the high occupancy sites (top ∼20%), and the fraction of ‘Very slow’ sites tends to decline with increasing occupancy.

**Figure 7.**
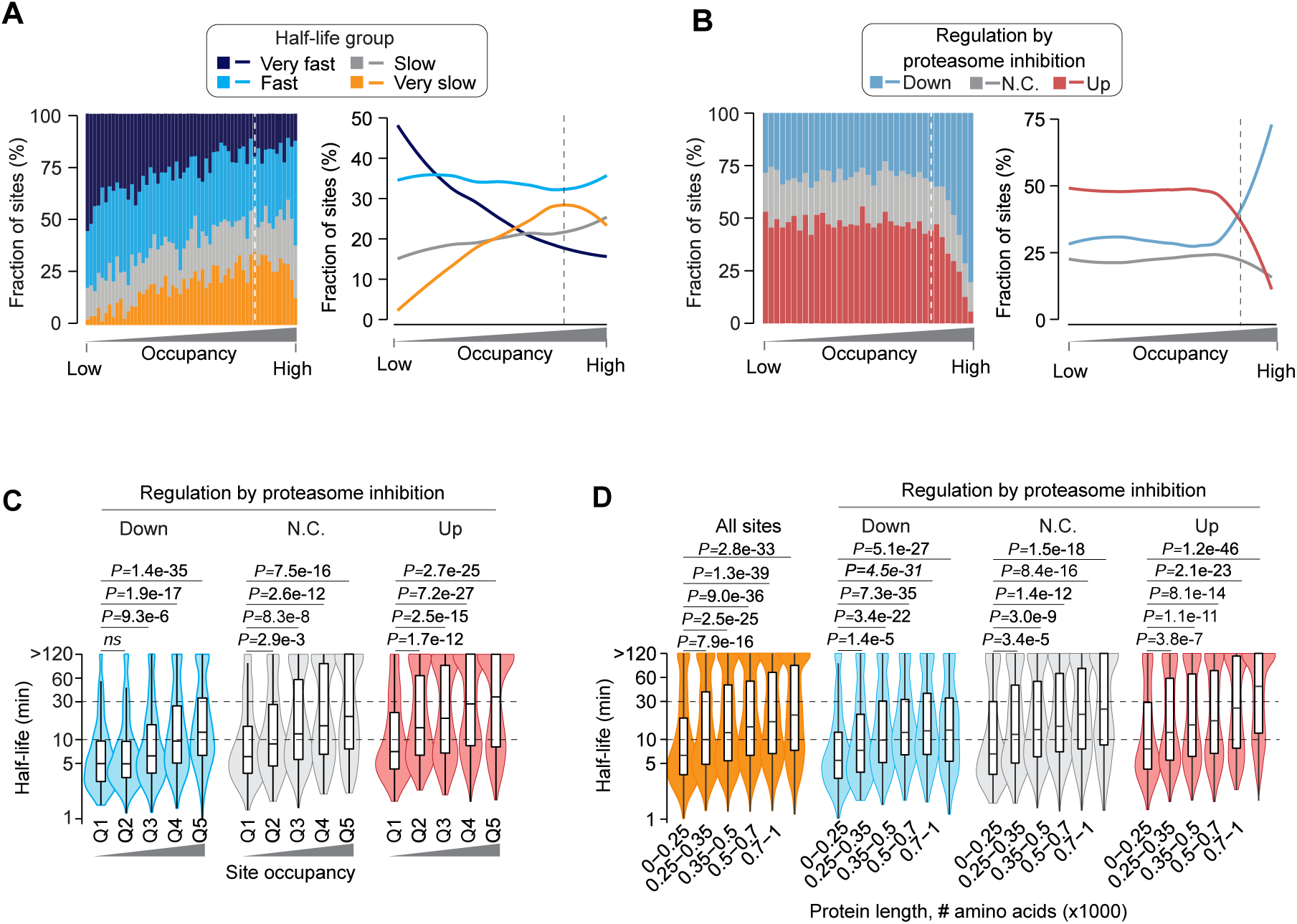
Strong interrelationships among ubiquitylation site occupancy, half-life, proteasome inhibition-induced changes, and protein length. (**A-B**) The global relationship between ubiquitylation site occupancy with the half-life (**A**), or between ubiquitylation site occupancy and proteasome inhibitor-induced ubiquitylation changes (**B**). UB sites were rank-ordered and binned based on their occupancy. Within each occupancy bin, the stacked bar charts (left panels) show the fraction of sites corresponding to the indicated half-life groups (**A**), or proteasome inhibitor-regulated class (**B**). The line charts (right, next to the bar charts) show global trends emerging from the shown bar charts. The global trends were determined by the local regression algorithm. The dotted line separates the lowest 80% and highest 20% occupancy sites. (**C-D**) The sites strongly upregulated by proteasome inhibitors, sites with high occupancy, and sites occurring in longer proteins have longer half-lives. First, sites were grouped based on their regulation after proteasome inhibition (Up: upregulated, Down: downregulated, N.C.: Not changed). Then, sites were further divided into quintiles based on their occupancy (**C**) or grouped based on the length of ubiquitylated proteins (**D**), as indicated. Box represents the interquartile range, the middle line indicates the median and the whiskers indicate the minimum and maximum values excluding outliers. P-values were calculated using a two-sided Wilcoxon rank-sum test (*ns*, P >0.05).

Next, we delved into the relationship between UB site occupancy and its regulation by proteasome inhibition. We ranked sites based on their occupancy and analyzed the fraction of sites regulated by proteasome inhibitors within each occupancy bin. For the bottom ∼80% of sites with the lowest occupancy, the fraction of both up- and down-regulated sites remained relatively constant (**Figure 7B**), despite the linear increase in half-life with occupancy (**Figure 7C**). However, an intriguing finding emerged when examining the highest occupancy sites. In the top ∼20% of sites, as occupancy increased, the fraction of upregulated sites decreased, while the fraction of downregulated sites sharply increased (**Figure 7B**). Among the bottom 100 sites with the lowest occupancy, 68% were upregulated by MG-132, whereas among the top 100 sites with the highest occupancy, only one site was upregulated, which corresponded to the K48 linkage of ubiquitin known for its involvement in proteasomal degradation (**Figure S7D**). This striking difference indicates that high occupancy sites are predominantly associated with non-proteasomal functions.

To further investigate these relationships, we categorized proteasome inhibitor-regulated sites into three classes: upregulated, downregulated, and not changing. Within each class, sites were grouped into occupancy quintiles. Across all classes, half-life increased with increasing occupancy, but the increase was most pronounced for the proteasome inhibitor upregulated sites (**Figure 7C**). This implies that sites highly upregulated by proteasome inhibitors are not necessarily turned over rapidly; rather, they exhibit slower turnover rates. Intrigued by this finding, we explored the association between UB site half-life and protein length. Surprisingly, a significant correlation between half-life and protein length was observed, with sites occurring in longer proteins displaying longer half-lives. This trend was most evident for sites upregulated after proteasome inhibition (**Figure 7D**). The underlying reasons for this association are unclear, but it is possible that longer proteins require more time to be targeted for proteasomal degradation, or the proteasome itself requires more time to efficiently degrade them, or the combination thereof.

Collectively, these results unveil intricate connections between UB site occupancy, half-life, and regulation by the proteasome. The fact that the measurements of occupancy, half-life, and proteasome inhibitor-induced ubiquitylation were performed entirely independently, the noteworthy relationships among these measurements strongly support the accuracy of our measurements.

## Discussion

The absolute abundance and turnover rate are crucial attributes of biomolecules, directly influencing their function. However, until now, comprehensive quantification of the absolute abundance and half-life has been limited to RNAs and proteins. This effort presents the most comprehensive quantification to date of the site-specific occupancy and half-life of a major eukaryotic PTM, providing insights into the abundance and turnover rates of ubiquitin-modified proteoforms.

Our comprehensive analyses build upon this qualitative understanding and provide a deeper quantitative comprehension of global ubiquitylation. To highlight the conceptual advance provided by our findings, we will focus on the emergent properties of the ubiquitylation system and their significance in understanding cellular dynamics. While some of these global properties may be speculated in retrospect, we are not aware of any study that has quantitatively predicted the ubiquitylation attributes revealed through our comprehensive systems-scale measurements. Here, we present five key findings from our work, which, to the best of our knowledge, have not been previously reported.

### 1. Site occupancies of different PTMs are vastly distinct

Ubiquitylation and phosphorylation have comparably broad regulatory scope, and the occupancy of ubiquitylation spans over 4 orders of magnitude, yet the global occupancy of ubiquitylation is more than 3 orders of magnitude lower than phosphorylation. We rationalize this observation as follows. A cell has a finite number of ubiquitin molecules, and ∼80% of ubiquitin molecules are already conjugated to proteins in cultured mammalian cells (Kaiser et al., 2011). Thus, even if all the remaining free ubiquitin is conjugated to proteins, it would only modestly increase in the median ubiquitylation occupancy. The limited amount of ubiquitin molecules presents an inherent bottleneck that restricts the site occupancy. On the other hand, the occupancy of phosphorylation is not constrained by the availability of ATP, allowing proteins to be comprehensively phosphorylated when necessary. For instance, during mitosis, the median phosphorylation occupancy reaches ∼70% (Olsen et al., 2010).

Achieving high global occupancy for ubiquitylation is impossible due to the inherent limitations. However, high occupancy can be achieved for specific ubiquitylation events if they target a limited number of sites or occur temporarily with rapid turnover. Interestingly, within this low occupancy regime, the occupancy of individual ubiquitylation sites varies by at least four orders of magnitude, which is comparable to the abundance range of mRNAs in mammalian cells (Schwanhüusser et al., 2011). We propose that low occupancy is a defining characteristic of ubiquitylation and may also apply to sumoylation, where the abundance of the SUMO protein is also limited. From the signaling perspective, ubiquitylation presents a paradigm for understanding how a limited quantity of signaling molecules can generate extensive complexity and regulate vast biological networks. Our finding revises the perception that low occupancy is biologically unimportant and underscores the need for precise quantitative measurements.

We note that the low site occupancy measured in our analyses sharply contrasts with previous study that quantified ∼150 sites and reported a median occupancy of 6%, with one third of sites having occupancy >20% (Hristova et al., 2020). Such high global ubiquitylation occupancies are theoretically improbable, and likely reflects inherent quantification limits of the method used for occupancy measurement (Prus et al., 2019). This emphasizes the difficulty in predicting ubiquitylation site occupancy or measuring it using standard mass spectrometry methods.

### 2. A wide dynamic range of ubiquitylation site kinetics

As such ubiquitylation is widely known as a dynamically regulated modification. Our work a offers a new, site-resolved view of ubiquitylation dynamics. To the best of our knowledge, this represents the first and largest analysis of site-specific turnover rates for any eukaryotic PTM on a systems scale. Our findings unveil that ubiquitylation, as a whole, exhibits turnover rates that are ∼50-200 times faster than those of mRNA and proteins. However, there is significant variability in the half-lives of specific sites, with individual site half-lives varying up to 100-fold. The rapid turnover of ubiquitylation help us rationalize how a small amount of ubiquitin can effectively regulate thousands of proteins. We conceive that, from the functional perspective, a low abundance of a site can be compensated by rapid flux such that sites with different occupancies can impact a similar number of protein molecules within a given time frame. Importantly, however, a site with low occupancy cannot simultaneously regulate as many protein molecules as a site with high occupancy.

### 3. Site occupancy and kinetics inform general properties and offer mechanistic insights

PTMs can either activate or inactivate enzymes, influence the subcellular localization of proteins, modulate protein-protein interactions (PPIs), or even affect the occurrence of other PTMs through a phenomenon known as ‘PTM cross-talk’. Our measurements of site occupancy suggest significant differences in how different PTMs mechanistically impact protein regulation. For instance, with broad and high site occupancies, phosphorylation possesses the potential to mediate both transient and long-term PPIs. In contrast, ubiquitylation, with its very low site occupancy, cannot facilitate high stoichiometry PPIs. Instead, it likely mainly promotes transient sub-stoichiometric interactions. Another implication of our finding is the extent to which a PTM can simultaneously impact the proteome. With its high global occupancy, phosphorylation is capable of simultaneously regulating a substantial portion of the proteome. This is exemplified by the dramatic increase in phosphorylation occupancy during mitosis, suggesting its role in mass-scale regulation (Olsen et al., 2010). On the other hand, ubiquitylation, with its very low site occupancy, cannot achieve such simultaneous, mass-scale regulation.

Considering that lysine residues are targeted by multiple PTMs, ubiquitylation can theoretically hinder the occurrence of other PTMs at the same lysines through competitive PTM cross-talk (Hunter, 2007). However, the low occupancy of ubiquitylation implies that over 99.9% of lysine residues remain non-ubiquitylated, suggesting that the involvement of ubiquitylation in competitive PTM cross-talk is likely limited.

In addition to uncovering general properties of ubiquitylation, our data reveal protein class-specific traits. We observed that plasma membrane-associated SLC proteins, as a class, have distinctively high occupancy sites in their cytoplasmic domains. Ubiquitylation plays a crucial role in regulating the endocytosis, trafficking, and sorting of various transmembrane receptors (Clague et al., 2012; Foot et al., 2017). Notably, ubiquitylation also controls the trafficking of membrane permeases in budding yeast (Lauwers et al., 2010), suggesting that the role of ubiquitylation in regulating nutrient transporter trafficking is anciently conserved.

Our site-resolves analyses can enable the inference of in vivo dynamics for processes regulated by individual ubiquitylation sites and contribute to the construction of mechanistic models. For example, the function of RQC is well established, but the frequency of RQC occurrence in a mammalian cell and the time required to rescue stalled ribosomes remain unclear. Considering that the number of ribosomes (3-6 × 10^6^) a HeLa cell (Bekker-Jensen et al., 2017; Duncan and Hershey, 1983), and the occupancy (median 0.29%) of RQC-associated sites, we can estimate that tens of thousands of ribosomes are ubiquitylated in steady-state Hela. If we consider RQC-associated sites as diagnostic markers of stalled ribosomes, this suggests that ribosome stalling is not a rare event; rather, it occurs frequently and potentially affects thousands of mRNA molecules at any given time. Furthermore, assuming that the kinetics of RQC site deubiquitylation reflects the resolution of stalled ribosomes, the rapid half-life (median ∼5 minutes) of RQC sites indicates that stalled disomes are rescued within minutes of signal initiation. We anticipate that our dataset will serve as a valuable resource for distinguishing a small number of signaling-relevant ubiquitylation sites from the vast number of sites involved in proteasomal degradation and will contribute to improving our mechanistic understanding of regulatory ubiquitylation sites.

### 4. A novel E1-E2 surveillance mechanism and its bypass by a bacterium

Ubiquitylation plays a critical role in regulating protein quality control by targeting mis-folded, unfolded, and fully folded proteins for degradation. We made an unexpected discovery of an ultra-rapid and site-indiscriminate deubiquitylation of all ubiquitin-conjugating E1-E2s. This raises the question of how E1-E2s become ubiquitylated and why they are deubiquitylated so quickly. While some E2s can autoubiquitylate (Banerjee et al., 1993; David et al., 2010), this is unlikely to be the case for E1s. Because fewer E1-E2s must collaborate with many more E3s, it is plausible that E3s, in addition to ubiquitylating their intended substrates, inadvertently ubiquitylate E1-E2s. This unintentional ubiquitylation could impair the function of E1-E2s, as previous studies have demonstrated that ubiquitylation can reduce the activity of E2s (Banka et al., 2015; Liess et al., 2019; Machida et al., 2006; McKenna et al., 2001; W. et al., 2005). An example is UBE2N, which is the most abundant E2 enzyme in cells (Clague et al., 2015b) and catalyzes most of K63 ubiquitin linkages in the cell. The activity of UBE2N is inhibited by modification at UBE2N^K92^ (T. and Yokosawa, 2005). Our analysis reveals that UBE2N^K92^ has an occupancy of 1.47% and a half-life of 1.6 minutes. If the modification of UBE2N^K92^ is not promptly removed, a rapid increase in occupancy can cripple its function.

We propose that the promiscuity of E3 enzymes, necessary for the rapid ubiquitylation of diverse proteins, unintentionally leads to collateral damage on E1-E2 enzymes. To counteract this unintended consequence, cells have evolved an elaborate and previously unappreciated surveillance mechanism. We suggest that this surveillance mechanism engages yet-to-be-identified deubiquitylases (DUBs) that remove bystander ubiquitylation on E1-E2 enzymes and mitigate the disadvantages associated with E3 promiscuity.

The importance of this surveillance mechanism is highlighted by the bacterium *Legionella pneumophila*, which appears to have evolved a way to bypass this mechanism. During early infection stages, the *L. pneumophila* effector MavC catalyzes a non-canonical type of ubiquitylation on UBE2N, where Q40 of ubiquitin is conjugated to UBE2N^K92^ through Nɛ-(ϒ-glutamyl)lysine isopeptide bond (Gan et al., 2019). Through this non-canonical ubiquitylation the bacterium inactivates UBE2N. In later infection stages, when the bacterium may require UBE2N function, the non-canonical ubiquitylation is removed by the bacterial deamidase MvcA (Gan et al., 2020). Until now, the reason why *L. pneumophila* employs non-canonical ubiquitylation for UBE2N inactivation had remained puzzling. Our finding, that canonical UBE2N^K92^ ubiquitylation is rapidly removed by the E1-E2 surveillance mechanism, suggests that *L. pneumophila* has evolved non-canonical ubiquitylation to circumvent this surveillance mechanism.

### 5. Ubiquitylation site occupancy and half-life are interrelated

Our investigation into the interrelationships between ubiquitylation occupancy, half-life, and the impact of proteasome inhibitors has uncovered notable distinctions between the properties of the lowest 80% and highest 20% occupancy sites (**Fig. 5 A-B**). In the bottom 80% of sites, we observe a nearly linear relationship between occupancy and half-life, while the fraction of sites regulated by proteasome activity remains relatively constant. In contrast, the top 20% of sites with the highest occupancy exhibit a decrease in the fraction of sites upregulated by proteasome inhibitors, accompanied by a sharp increase in the fraction of downregulated sites. This suggests that the differences in ubiquitylation site occupancy arise from two distinct phenomena. Among the bottom 80% of sites with the lowest occupancy, the differences in half-life, at least partially, account for the variation in occupancy observed. However, for the top 20% of sites with the highest occupancy, their high levels cannot be solely explained by half-life, strongly suggesting that the high occupancy of these sites results from targeted ubiquitylation by site-specific ligases.

In summary, our findings reveal the general properties of ubiquitylation and uncover the design principles of ubiquitylation-dependent cellular governance. It appears that ubiquitin functions akin to a currency in the cellular signaling system, where a limited amount of this ‘currency’ is strategically allocated across thousands of proteins and tens of thousands of sites. The ubiquitylation economy is designed to operate efficiently with its inherent constraints of limited availability and rapid turnover, much like the role of monetary currency in the global economy.

#### Study limitations

We examined the properties of ubiquitylation in cultured cells under standard laboratory conditions. The occupancy of sites may vary in different conditions and cell types. However, it is reasonable to assume that the general characteristics of ubiquitylation are conserved across different systems.

Our work does not inform on the specific function or regulatory mechanisms of individual ubiquitylation sites. Our primary focus was to uncover the fundamental design principles of ubiquitylation by quantifying global site occupancy and half-life. Rather than investigating the biological function of any particular site, protein, or pathway, we aimed to provide a broad understanding of the overall ubiquitylation process.

The presence of distinctly rapid and indiscriminate deubiquitylation of E1-E2 enzymes strongly suggests the existence of a dedicated mechanism for their rapid deubiquitylation. However, the precise identity of the specific DUBs involved in this process remains to be determined and requires further investigation.

## Acknowledgments

We thank members of the Choudhary laboratory for their helpful discussions. This project received funding from the European Research Council (ERC) under the European Union’s Horizon 2020 research and innovation program (grant agreement: 648039). GP is supported by the Novo Nordisk Foundation (grant agreement: NNF17CC0026748). We thank the Big Data Management and Mass Spectrometry Platforms at CPR for their excellent support. The Novo Nordisk Foundation Center for Protein Research is supported financially by the Novo Nordisk Foundation (Grant agreement: NNF14CC0001).

## Author contributions

B.T.W. and S.S. developed the method for quantifying ubiquitylation occupancy. G.P. performed most of the experimental work and analyzed the data. T.N. helped G.P. in the calculation of ubiquitylation occupancy. C.C. and G.P. wrote the manuscript, with input from all authors. C.C. conceived and supervised the project.

## Inclusion and diversity

We support inclusive, diverse, and equitable conduct of research.

## STAR★Methods

### Resource availability

#### Lead contact

Further information and requests for resources and reagents should be directed to and will be fulfilled by the lead contact, Chunaram Choudhary.

#### Experimental model and study participant details

HeLa cell line was purchased from American Type Culture Collection (ATCC) and cultured in DMEM media (BioWest) supplemented with 10% dialyzed fetal bovine serum (FBS, Sigma-Aldrich), 2 mM L-Glutamine (Thermo Fisher Scientific), 50 units/mL of penicillin, 50 µg/mL of streptomycin (Roche). For SILAC labeling, cells were cultured in SILAC media for at least eight passages containing either unlabeled ’light’ (L) amino acids (L-lysine; L-arginine), ’medium’ (M) amino acids (D^4^ L-lysine, 13C^6^ L-arginine), or ’heavy’ (H) amino acids (13C^6^ 15N^2^ L-lysine, 13C^6^ 15N^4^ L-arginine) (Cambridge Isotope Laboratories) as previously reported (Ong et al., 2002). Cells were grown in a 37^0^C humidifier incubator and were passaged upon attaining 70-80% confluence.

### Method details

#### Preparation of inhibitor stocks and storage

Cells were treated with the ubiquitin-specific E1 inhibitor TAK-243 (also known as MLN7243, Chemgood), and proteasome inhibitors: MG-132 (Enzo Life Sciences), and Bortezomib (BTZ; Selleckchem). Unless indicated otherwise, cells were treated with a final concentration of 100µM TAK-243, 200µM MG-132, and 100µM BTZ.

#### Ubiquitylation site occupancy measurements

##### Cell lysis

Before harvest, cells were washed twice with PBS and detached by trypsinization at 37°C. The cells were centrifuged at 500g for 5 minutes, and the pellet was washed twice with ice-cold PBS, followed by snap-freezing in liquid nitrogen. Before preparing samples for mass spectrometry analyses, cells were lysed in 6M Guanidium hydrochloride (GndHCl), 50mM HEPES pH 8.0, 1mM TCEP, 5mM CAA, and the sample was sonicated (Branson Ultrasonics) for 30 seconds at 70% power. To maximize protein extraction, and to ensure the inactivation of proteases, especially deubiquitinases (DUBs), proteins were denatured by incubating the lysates at 95°C (10 min). The lysates were cleared by centrifugation at 13,000 × g for 30 minutes. Proteins were quantified by Quick Start^TM^ Bradford assay (BioRad) and protein concentration was adjusted to 6 mg/ml using 100mM HEPES pH8.

##### Preparation of partial chemical GG-modified spike- in standard

To generate a partially chemically GG-modified spike-in standard, heavy SILAC-labeled HeLa cell lysates were incubated with NHS-Gly-Gly-Boc ester (NGGB; A ChemTek Inc.). The proteins were treated with 1mM NHS-Gly-Gly-Boc ester at 30°C for 1 hour. We prepared two independent batches of partially chemically GG-modified protein (PC-GG) standard PC-GG. To determine the degree of PC-GGPC-GG, we generated a comprehensively chemically modified protein (CC-GG) reference. For this, light SILAC-labeled proteins were incubated overnight with 100mM NHS-Gly-Gly-Boc ester at 37°C. The reaction was quenched by adding Tris-HCl pH 8.0 to a final concentration of 250mM.

The degree of partial chemical GG modification was quantified by spiking in comprehensively chemically GG-modified proteins. The light SILAC-labelled CC-GG proteins were mixed with heavy SILAC-labeled PC-GG proteins at three different dilutions (1:10, 1:50, 1:250) to yield GG site occupancy of 10%, 2%, and 0.4%. The sample mixtures were digested, desalted, and analyzed on a mass spectrometer.

Because the lysine modification prevents cleavage by trypsin, chemical modification of lysine causes a proportional reduction in the abundance of the corresponding fully cleaved unmodified peptides (Weinert et al., 2017). Therefore, we quantified the reduction in peptides cleaved at lysine to independently verify the degree of PC-GG. We mixed a heavy-labeled PC-GG sample with unmodified light-labeled proteins in a 1:1 ratio. The protein mixture was digested with trypsin overnight, the resulting peptides were desalted and analyzed on a mass spectrometer.

##### Sample preparation for native ubiquitylation occupancy measurements

To determine native ubiquitin occupancy, SILAC-heavy-labeled PC-GG was spiked-in into native SILAC-light-labeled protein lysate at different dilutions to yield a ubiquitylation occupancy of ∼3%, ∼1%, ∼0.1%, ∼0.01%, and ∼0.001%. Before proteolysis, the concentration of GndHCl in lysates was reduced to 1M by diluting the samples with 50mM HEPES pH 8.0, and the proteins were digested overnight with trypsin (enzyme to protein ratio 1:100; weight/weight). Trypsin proteolysis was stopped by adding 1% TFA, and the resulting peptides were desalted on Sep-Pak C18 cartridges (Waters). Briefly, before sample loading, C18 material was activated with 100% acetonitrile (ACN) and equilibrated with 0.1% trifluoroacetic acid (TFA). Acidified peptides were loaded and washed with 0.1% TFA. Peptides were eluted with 50% ACN 0.1% TFA and were vacuum centrifuged to evaporate ACN. To remove the BOC protective group from PC-GG-modified peptides, the purified peptides were incubated with 33% TFA for 1 hour. The TFA solution was diluted to 1.5%, and the peptides were re-purified by Sep-Pak C18 cartridges.

##### Sample preparation for ubiquitylation half-life quantification

To quantify ubiquitylation half-life, the cells were treated with the E1 inhibitor TAK-243 (100µM) for 0, 5, 10, 30, and 60 min. To cover all the time points, we performed two sets of triple SILAC experiments. In set 1, medium or heavy-labeled cells were treated with TAK-243 for 5 and 30 minutes, respectively. In set 2, medium or heavy-labeled cells were treated with TAK-243 for 10 and 60 minutes, respectively. In both sets, mock-treated light SILAC-labeled cells were used as control. Cells were quickly washed with PBS and lysed on the plate in the sodium deoxycholate (SDC) buffer (2% SDC, 50mM HEPES pH 8.0 supplemented with 5mM NEM). The lysates were collected and immediately boiled at 95°C for 5 min to denature proteins The samples were placed on ice and sonicated (Branson Ultrasonics) for 1 minute at 70% amplitude. The lysates were cleared by centrifugation at 4.000g for 15 minutes. Protein concentration was estimated using the BCA Protein Assay kit (Thermo Fisher Scientific), and the samples from SILAC light-, medium-, and heavy-labeled cells were mixed in a 1:1:1 ratio. The mixed samples were treated with DTT (7.5mM, at 70oC for 10 minutes) to reduce disulfide bonds, and the reaction was quenched by adding NEM (10mM). The samples were proteolyzed overnight (37°C) with trypsin (enzyme to protein ratio 1:100 (w/w)). To stop the reaction and remove SDC from the sample, we added TFA to the final concentration of 1%. The peptide mixture was subsequently cleared by centrifugation at 4.500g for 30 minutes. The peptides were desalted using SepPak C18 cartridges as described above.

##### Sample preparation for ubiquitylation analysis after proteasome inhibitor treatment

To investigate the role of proteasome activity in ubiquitylation turnover, cells were treated either with the proteasome inhibitors MG-132 (200µM) or Bortezomib (BTZ, 100µM), alone, or in combination with the UAE1 inhibitor TAK-243 (100µM). In one setup, we analyzed the effect of E1 inhibition alone or the simultaneous inhibition of E1 and proteasome. For this, SILAC-labelled (light, medium, heavy) HeLa cells were treated for 1h either with DMSO, TAK-243, or TAK-243 in combination with MG-132. In another setup, we analyzed the effect of proteasome inhibition alone or sequential inhibition of the proteasome and E1. In this setup, SILAC-labelled (light, medium, heavy) cells were treated for 2.5 hours with DMSO, the proteasome inhibitors (BTZ, or MG-132), or with the proteasome inhibitors (BTZ or MG-132) and E1 inhibitor (TAK-243), where E1 inhibitor was administrated 30 minutes before harvest. The samples were processed as described above for quantification of the UB site half-life.

##### GG-modified peptide enrichment and sample fractionation

To reduce the sample complexity and to increase the depth of GG remnant profiling, the samples were fractionated using high-pH reversed-phase (HpH) alone or in combination with strong cation exchange (SCX) chromatography. The samples were fractionated before or after GG-modified peptide enrichment.

For fractionating peptides by HpH before di-Gy enrichment, ∼15mg of peptides were dissolved in 20mM ammonium hydroxide and fractionated using a Dionex Ultimate 3000 HPLC as described previously (Batth et al., 2014). A total of 24 fractions were collected and concatenated into six fractions, followed by vacuum centrifugation to dryness. Each of the 6 fractions was dissolved in 100µl 10x the immune affinity purification (IAP) buffer (500 mM MOPS; pH 7.2, 100 mM Na-phosphate, 500mM NaCl, 2.5% NP-40) and diluted to a final concentration of 1x IAP in 1ml. We used PTMScan® Ubiquitin Remnant Motif (K-ε-GG) antibody kit (Cell Signalling Technology) for the enrichment of GG-modified peptides. A single tube of antibody agarose beads (80µL) was equally split into six equal portions and mixed with GG-modified peptides from 6 HpH fractions. Samples were incubated with beads on a rotating wheel for 2 hours at 4°C. The beads were washed two times each with IAP buffer, IAP buffer without NP40, and deionized water. Enriched peptides were eluted with 0.15% TFA. Each fraction was further fractionated into three fractions using the micro-SCX state-tip format, as described previously (Weinert et al., 2013) with minor changes. Briefly, SCX material (3M, Bracknell) was activated with 100% methanol and equilibrated with 50% ACN 0.1% TFA. After sample loading, the column was washed with 100µl of 50% ACN 0.1% TFA. Peptides were eluted in elution buffer (20mM acetic acid, 20mM boric acid, 20mM phosphoric acid, 50% ACN), gradually increasing pH 4.0-9.0.

For fractionating peptides after GG-modified peptide enrichment, approximately 15 mg of SepPak C18 desalted peptides were reconstituted in IAP buffer, mixed with pre-washed anti-K-ε-GG antibody-conjugated beads (80 µl), and incubated at 4°C for 1 hour. The beads were washed two times each with IAP buffer, IAP buffer without NP40, and deionized water.

Enriched peptides were eluted with 0.15% TFA. To remove any remaining detergents (e.g. NP-40) in the sample, the sample was further cleaned up using SCX Stagetips. Peptides were eluted with the high pH (pH 9.0) elution buffer, and the organic solvent was removed by vacuum centrifugation. We used a self-made tip-based column for HpH fractionation of enriched GG-modified peptides where 3μm ReproSil-Pur 120A C18-AQ beads (Dr. Maisch GmbH) were trapped on top of a single C8 Empore disc membrane (3M, Bracknell, UK). After bead activation and equilibration, peptides were loaded and separated at pH 10.0 with 10mM ammonia formate as a mobile phase. Peptides were eluted with an increasing gradient of ACN (8, 10, 12, 14, 16, 18, 20, 22, 24, and 50%).

##### Sample preparation for total proteome measurements

For all GG remnant measurement experiments with E1 and proteasome inhibitors, we also measured the proteome to correct for SILAC mixing. Briefly, before GG-modified peptide enrichment, we took out a small fraction (∼50ug) of peptides and fractionated them into four fractions using in-tip HpH fractionation, as described above. Peptides were eluted with an increasing gradient of ACN (10, 16, 20, and 50%), and analyzed by MS.

##### LC-MS/MS analysis

Before MS analysis, peptides were desalted on a C18 homemade Stage-tip. Peptides were analyzed by nanoflow liquid chromatography coupled to tandem mass spectrometry (nLC-MS/MS). The sample was loaded on a 15cm capillary column packed with 1.9μm Reprosil-Pur C18 beads (Dr. Maisch) attached to an EASY-nLC 1000 system (Thermo Scientific).

Peptides were separated with an increasing gradient of ACN in 0.1% FA at 300nL/min flow rate. The column temperature was maintained stable at 50°C using an integrated column oven (PRSO-V1, Sonation GmbH). Eluting peptides were analyzed in a positive mode in data-dependent acquisition mode by either Q-Exactive HF, Q-Exactive HF-X, or Exploris 480 (Thermo Scientific). We used different acquisition parameters (summarised in Table 1) for each instrument to maximize the performance. Q-Exactive HF and Q-Exactive HF-X were operated with Tune 2.3 and Xcalibur 3.0.63. Exploris 480 was operated with Tune and Xcalibur 4.4.

**Table 1.** Acquisition parameters for mass spectrometry analysis.

|  | Q-Exactive HF | Q-Exactive HF-X | Exploris 480 |
| --- | --- | --- | --- |
| **Full Scan Settings:** |  |  |  |
| Resolution (at m/z 200) | 60 000 | 60 000 | 120 000 |
| Injection time (ms) | 20 | 20 | 60 |
| AGC | 3.0e6 | 3.0e6 | 300% |
| **DDA-MS2 Settings:** |  |  |  |
| Intensity threshold | 5.0e4 | 1.0e5 | 5.0e4 |
| Charge state | >1 | >1 | 2-5 |
| Dynamic exclusion (s) | 30 | 30 | 30 |
| Isolation window (m/z) | 1.3 | 1.3 | 1.3 |
| Resolution (at m/z 200) | 30 000 | 15 000 | 15 000 |
| Injection time (ms) | 250 | 110 | 110 |
| NCE (%) | 28 | 28 | 25 |
| AGC | 2.0 e5 | 1.0 e5 | 80% |

##### Estimation of total protein and ubiquitin content in HeLa

To determine total protein content in a HeLa cell, one million SILAC light-labeled HeLa cells were lysed in SDC lysis buffer (2% SDC, 50mM Hepes pH 8.0 supplemented with 5mM NEM), and boiled at 95°C for 5 min. The lysates were placed on ice, sonicated (Branson Ultrasonics) for 1 minute at 70% amplitude, and cleared by centrifugation at 4.000g for 15 minutes. Total protein concentration in HeLa cells was estimated using the BCA Protein Assay kit to 158±15 pg/cell.

To determine ubiquitin concentration in HeLa, light SILAC-labeled HeLa proteins were digested with trypsin, and peptides were desalted on the Sep-Pak C18 column, as described above. Heavy isotope-labeled synthetic ubiquitin peptide (ESTLHLVLR[+10Da]) was spiked-in into the peptide mixture in three 2-fold serial dilutions to yield a concentration of 50, 100, and 200fmol per 100ng of HeLa peptides. Samples were measured on Exploris 480 mass spectrometer (Thermo Scientific), coupled to EASY-nLC 1000 system (Thermo Scientific). Peptides were separated on a 50-minute acetonitrile gradient (2-40% ACN). Eluting peptides were analyzed using the parallel reaction monitoring (PRM) method (Peterson et al., 2012). Ions of 539.3182(2+), 359.8812(3+), 534.314(2+), and 356.5451(3+) m/z values were filtered with 1.3 m/z isolation window, maximum injection time of 100ms, normalized AGC target 80%, and fragmented with HCD collision energy set to 25%. MS2 spectra were collected at the 30.000 orbitrap resolution.

Raw files from PRM measurements were converted to mzXML format using MSConvert v3.0.18285-e65c323cc (ProteoWizard) and analyzed in Skyline software v20.1.0.155 (MacLean et al., 2010). Fragments spectra were compared to a reference spectral library generated by MaxQuant search of HeLa proteome. Fragments peaks were manually checked for the spectra library match, RT alignment, and peak shape. The Savitzky-Golay smoothening was applied. Any fragments suspected of the presence of interference from other ions were removed. To calculate the ubiquitin content in HeLa cells, the peak area of native peptides (light SILAC-labelled) was normalized to the level of heavy-labeled peptides and corrected for the synthetic peptide concertation. From two biological replicates, we estimated a ubiquitin concentration of 477±68 pmol/mg of HeLa proteins, which is similar to the previously reported ubiquitin concentration (486.4 ± 42 pmol/mg) in HEK293 cells (Kaiser et al., 2011). Using HeLa cell protein content of 158 pg/cell, we estimated 4.54×10^7^ ubiquitin molecules/cell in HeLa. It is estimated that a HeLa cell contains ∼4×10^9^ protein molecules/cell (Bekker-Jensen et al., 2017), therefore, we estimate that ubiquitin constitutes 1.1% of protein molecules.

##### Immunoblot analysis

Samples were denatured in NuPAGE LDS Sample Buffer (Invitrogen, Thermo Fischer Scientific) with the addition of 100mM DTT at 95°C for 5 minutes. Proteins (20µg) were separated on 4-12% NuPAGE gel (Invitrogen, Thermo Fischer Scientific) and transferred onto a nitrocellulose membrane (BioRad). After blocking with 5% BSA (Sigma-Aldrich) in PBST (PBS, 1% Tween), membranes were incubated overnight with primary antibodies at 1:1000 dilution at 4°C. Membranes were washed with PBST, incubated with horseradish peroxidase (HRP)-conjugated secondary antibodies (1:10000 dilution) for 1 hour at 4°C, and washed again with PBST. Proteins were visualized using ECL Chemiluminescent Substrate Reagent Kit (Novex, Thermo Fisher Scientific). The same membrane was stripped off (0.5M glycine buffer pH 2.5) and re-blotted with an anti-β-actin antibody for loading control.

#### Data processing and analysis

##### RAW data processing

Peptide identification and quantification were performed using MaxQuant software (version 1.5.3.30) with the integrated Andromeda search engine (Cox and Mann, 2008). The data were searched against the human UniProt database (retrieved on April 26, 2018; containing 20,328 entries) with the following Andromeda settings: initial search mass tolerance of 20 ppm, main search mass tolerance of 6 ppm for parent ions, and 20 ppm for HCD fragment ions, trypsin specificity with a maximum of two missed cleavages, peptide length greater than five amino acids. Depending on the sample preparation buffers used, cysteine carbamidomethylation (+57.021 Da) and cysteine modification with NEM (+125.048 Da) were included as fixed modifications, and N-acetylation of proteins (+42.011 Da), deamidation on asparagine or glutamine (+0.984 Da), oxidized methionine (+15.995 Da), and GG-modified lysine (+114.043 Da) were included as variable modifications. Except for the occupancy measurements, the re-quantify option was enabled for quantifying GG-modified sites in all other experiments.

##### Estimation of the degree of partial chemical GG modification (PC-GG)

We generated two batches of PC-GG proteins. We assume that chemical modification is non-site-selective and all lysines are modified to a similar extent in denatured proteins. To estimate the degree of partial chemical GG-modification, we mixed heavy SILAC-labeled PC-GG proteins with light SILAC-labeled comprehensively chemically GG-modified proteins in the ratio of 1:10, 1:50, and 1:250. Following trypsin proteolysis, peptides were purified on C18 StageTips and analyzed by MS. The degree of PC-GG was calculated based on the median heavy-to-light SILAC ratio of GG-modified peptides, corrected for the CC-GG dilution (**Figure S1B**). We only used fully tryptic GG-modified peptides (without miscleavage, except for miscleavage caused by GG modification of lysine) that were generated by cleavage at arginine. The reason for using only arginine-cleaved peptides is that their trypsin digestion is not expected to be affected by the GG modification of lysine. To increase the quantification accuracy, we removed peptide intensity ratios that did not follow a serial dilution, allowing for up to two-fold variability between the dilutions. Based on this, we estimated the degree of PC-GG to be 4.04% for batch 1 and 11.28% for batch 2 of PC-GG-modified proteins (**Figure S2 A-B**).

Chemical GG modification of lysine prevents trypsin cleavage at the modified lysine and causes a decrease in the abundance of the corresponding unmodified peptides, and the decrease is proportional to the degree of chemical modification (Weinert et al., 2017). Therefore, we quantified the decrease in corresponding peptides to corroborate the degree of PC-GG estimated by spiking in CC-GG proteins. We mixed CC-GG modified (1:1) proteins (SILAC heavy) with proteins from control cells (SILAC light). Firstly, we classified fully-cleaved peptides based on their tryptic cleavage pattern: K0 peptides contain no lysine and therefore their abundance is not expected to be affected by the lysine modification, K1 peptides are generated by the cleavage at one lysine, and K2 peptides were flanked by the two lysines. For each of the peptide types, we separately quantified the median heavy-to-light SILAC ratio to determine the reduction in peptide intensity in the PC-GG sample. Because K0 peptides are not affected by the lysine modification, median SILAC ratios for K1 and K2 peptides were normalized by the median SILAC ratio of K0 peptides. Trypsin cleavage of K2 peptides is twice more likely to be affected by the modification compared to K1 peptides. Thus, the difference between the normalized median ratio of K1-K0 peptides, and K2-K1 peptides reflects the degree of PC-GG (**Figure S2C**).

##### Estimation of native ubiquitination site occupancy

To calculate site occupancy, we used SILAC ratios of singly modified peptides from the “evidence.txt” MaxQuant output table. We did not use the “GlyGly (K)Sites.txt” file because the SILAC ratios of individual sites can be derived from several different peptides. Likewise, individual entries in the “modificationSpecificPeptides.txt” table can include different positions of GlyGly within the same peptide sequence. Starting with the ’evidence.txt’ table, the following actions were performed:

1. Data were filtered for reverse decoy database entries, potential contaminants, and peptides with GlyGly (K) localization probability less than 75% (0.75.) Only singly GlyGly(K)-modified peptides were analyzed.
2. The “Modified sequence” was used as a unique identifier to calculate the median SILAC ratio and summed peptide intensity.
3. SILAC ratios were adjusted to account for the degree of PC-GG and the impact of the lysine modification on the peptide cleavage, for example, 10% PC-GG results in a 10% reduction in peptides generated by cleavage at a single lysine and a 20% reduction in peptides generated by cleaving two lysines (equation 1).

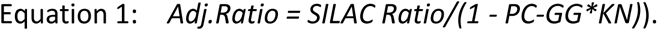

Where: *SILAC Ratio -* SILAC ratio between PC-GG and native GlyGly modified peptides, *PC-GG*- degree of PC-GG, KN-number of lysines involved in the digestion of a given peptide SILAC ratios were tested for agreement with the dilution series, allowing for up to two-fold variability, as previously described (Weinert et al., 2017). Occupancy was calculated using only peptides that met these strict criteria.
4. In each experiment, stoichiometry (occupancy) was calculated using equation 2.

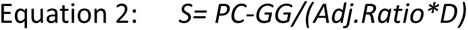

Where: *S* – stoichiometry (occupancy), *PC-GG* - degree of PC-GG, *D* - dilution factor for PC-GG peptides, *Adj.Ratio* - adjusted SILAC intensity ratio between PC-GG and native GlyGly modified peptides.

The dilution factors (*D*) were as follows: Experiment 1 - (3% *D*=1.25), (1% *D*=3.77), (0.1% *D*=37.7), (0.01% *D*=377), Experiment 2 - (1% *D*=11.3), (0.1% *D*=113), (0.01% *D*=1,130), (0.001% *D*=11,300), Experiments 3-6 - (1% *D*=10), (0.1% *D*=100), (0.01% *D*=1,000), (0.001% *D*=10,000).

##### Identification of phosphorylation and UB sites in deep proteome measurements

To identify the number of GG-modified peptides detectable in total proteome measurements, without GG-modified peptide affinity enrichment, we searched a previously published deep HeLa cell proteome dataset (Bekker-Jensen et al., 2017). To compare the detection frequency of phosphorylated and GG-modified peptides, the same dataset was simultaneously searched for phosphorylation. Raw files were downloaded from the PRIDE repository (PXD004452) and analyzed using MaxQuant (version 1.5.6.5) using the human proteome Uniprot database (retrieved on July 6, 2015) as a reference. The following search parameters were used: precursor and fragment ion mass tolerance of 20 ppm and 0.5Da, respectively, maximum of two cleavages, minimum length of peptides 7 amino acids, cysteine carbamidomethylation (+57.021 Da) was included as fixed modification, and protein N-term acetylation (+42.011 Da), methionine oxidation (+15.995 Da), serine, threonine and tyrosine phosphorylation (+79.966 Da) and internal lysine GG (+114.043 Da) were included as variable modifications. A default MaxQuant peptide identification score (40) and delta score (6) was applied for modified peptides. FDR was set to 1% at the peptide-spectrum match (PSM) and razor protein level. Following this “standard search,” the number of identified peptides (unmodified, phosphorylated, and GG-modified) was counted using the MaxQuant output table “modificationSpecificPeptides.txt,” after removing peptides originating from potential contaminants and decoy entries.

To remove low-confidence identifications, we filtered the data by applying the following criteria:

1. Peptides with a delta score <20 were removed.
2. The inclusion of multiple variable modifications increases the database search space and can result in a high false discovery rate (FDR) among multiply modified peptides, especially for low abundant modifications (Bogdanow et al., 2016). Therefore, peptides modified with more than one variable modifications (i.e phosphorylation, GG, or methionine oxidation) were removed.
3. MS-detected peptides must have at least one positive charge, and due to an additional amino terminus, GG-modified peptides are expected to contain an extra positive charge. We excluded GG-modified peptides with a charge state of +1. Unmodified and phosphorylated were not filtered based on their charge state.
4. Peptides identified as a second peptide in MS/MS fragment spectra were filtered out.

##### Validation of the UB site occupancy by abundance corrected intensity of GG-modified peptides (ACI)

To calculate abundance-corrected GG-modified peptide intensity (ACI), we divided the summed GG-modified peptide intensity with the iBAQ value of the corresponding protein.

Protein iBAQ values were calculated using the HeLa cell proteome dataset (Bekker-Jensen et al., 2017). Bekker-Jensen et al. used multiple enzymes and analyzed the samples using different types of MS instrumentation. For determining iBAQ values for our analyses, we only used peptides from samples digested with trypsin and analyzed them on Orbitrap instruments (these were the same settings used for analyzing GG-modified peptides in our analyses). Protein iBAQ values were calculated using 7-30 amino acid-long tryptic peptides.

##### Estimation of the correlation between MS intensity of individual peptides and protein iBAQ

We rationalized that peptides derived from the same protein must have equimolar abundance, and therefore, peptides derived from the same proteins could be used to determine the empirical correlation between the MS intensity of individual peptides and the iBAQ-based abundance of the corresponding protein. We chose all proteins that were identified with ≥5 peptides. From each protein, we arbitrarily selected one peptide and used the remaining peptides (minimum 4 peptides for each protein) to re-calculated iBAQ values of the corresponding protein. To ensure that randomly selected peptides were not biased, we performed three iterations, each time choosing a different random peptide. The correlation was determined between the MS intensity of arbitrarily selected peptides and the iBAQ values of their corresponding proteins.

##### Calculation of ubiquitination half-life

UB site half-life was calculated using the time-course data from TAK-243 treated cells. The MaxQuant output table ’GlyGly(K) sites’ was first filtered to remove reverse decoy database entries, potential contaminants, and sites with a localization probability below 90% (0.9).

For all GG remnant analyses, we also analyzed protein expression. To correct for any mixing errors, SILAC ratios of GG-modified peptides were corrected using the median SILAC ratio of the proteome measurement. We excluded outliers (<3% of data) and data with ratios equal to or above 1.5 (<2% of data), as we expected that after the inhibition of the ubiquitylation cascade, all UB sites have decreased or unaltered levels. The turnover rate was analyzed for sites where we obtained the quantitative data for all 4-time points (5, 10, 30, and 60 minutes). We performed a non-linear regression to fit an exponential function (equation 3) to our data.

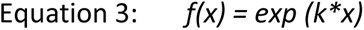

Where: *x* - time(min), *k* - decay constant,

While we obtained a good fit at early time points, we noticed that for many sites deubiquitylation reaches a plateau later for many sites (**Figure 2D**). We reasoned that this is likely because the ubiquitylation level is reduced below the linear quantification range in our analyses. To account for that, we applied an exponential plateau function (equation 4).

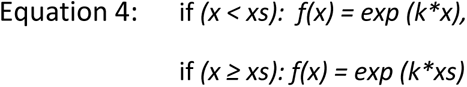

Where: *x* – time (min), *k* - decay constant, *xs* – time point at which plateau is reached.

For each UB site, we fit both exponential and exponential-plateau functions and the most suitable model was chosen based on Akaike’s information criterion (AIC) (Cavanaugh, 1997), which balances the goodness of fit and model flexibility. A low number of data points can lead to the overfitting of AIC, and therefore, we applied correction for the small sample size (Cavanaugh, 1997). For each model, the AIC was calculated as follows (equation 5):

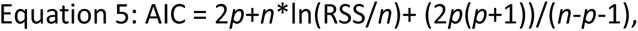

Where: *p* - number of parameters (for exponential model *p*=1 and for exponential-plateau function *p*=2), *n* - number of data points, RSS - root sum squares.

The model with a lower AIC value was chosen as the best fit.

UB site half-life was determined based on the range where the decay follows an exponential trend in the selected model (equation 6).

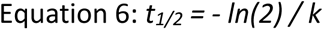

Where: *t_1/2_* - half-life (min), *k* - decay constant.

For a fraction of sites, ubiquitylation was not appreciably reduced within 1h or TAK-243 treatment. These sites were assigned a half-life >120 min (half-life at least twice as long as the longest measurement time point in our analyses, 60 min).

Based on their turnover rate, we grouped UB sites into ’Very fast’, ’Fast’, ’Slow’, and ’Very slow’, corresponding to a half-life of <5, 5-15, 15-60, >60 minutes. We used a *ComplexHeatmap* R package to generate a UB site half-life heatmap.

#### Gene Ontology term enrichment analysis

Gene Ontology (Biological Process, Molecular Function, and Cellular Compartment) enrichment was performed in R using *gprofiler* package (Kolberg et al., 2020). We converted Uniport IDs into Entrez Gene IDs, which were used as input. The enrichment analysis was performed against a user-defined background. For the enrichment analyses of proteins containing high occupancy (>0.5%) UB sites, all ubiquitylated proteins identified in the study were used as background reference. For the enrichment analysis of proteins harboring sites of different turnover rates, the background consisted of all proteins for which UB site half-life was calculated. Significantly enriched terms, with a Bonferroni p-value adjustment for multiple testing, are reported. We visualized selected, non-redundant Gene Ontology terms enriched in a high UB occupancy-containing subset of proteins with a p-adj<1.5%. To visualize enrichment in dynamically regulated Ub-proteins, we applied an additional filter and required that at least 60% of the proteins assigned to a GO term must be enriched. The full list of the enriched GO terms is included in **Table S4**. A protein may contain single or multiple sites with high occupancy, or turnover rates. Enrichment analyses at the protein level do not account for the number of sites within proteins. To account for that, for each term, we calculated the UB site enrichment (i.e., the fraction of all sites mapping to the enriched subset of proteins).

##### Classification of UB sites based on the regulation by proteasome inhibition

For all GG remnant measurement experiments, we took out a small portion of the total peptides and quantified total protein abundance in different SILAC states. Using the median SILAC ratio from the proteome measurements, we corrected SILAC ratios of GG-modified peptides to account for any SILAC mixing error. Sites showing ≥2-fold increased ubiquitylation after BTZ (100µM) or MG-132 (200µM) treatment (2.5 hours) were classified as upregulated (Up), sites showing ≥2-fold decreased ubiquitylation were classified as downregulated (Down), and sites showing less than 2-fold change were classified as not changing (N.C.) For sites quantified in both MG-132 and BTZ experiments, we applied additional prerequisites. UB site was classified as N.C. if ubiquitylation fold-change was between 1-2 for both the inhibitors. UB sites were classified as upregulated if their FC was >2 for at least one inhibitor and ≥1.5-fold in the other inhibitor. Similarly, sites were classified as downregulated if their FC was <1 in at least one inhibitor but not higher than ≤1.5 in the other inhibitor. A small number of sites (n= 289) did not match these criteria and were considered ambiguous and were not used for the analyses. This classification is used in further analysis to distinguish proteasome-dependent and -independent UB sites.

##### Mapping UB sites in individual proteins and domains

We classified transmembrane (TM) and plasma membrane (PM)-associated transmembrane proteins based on the following criteria. Proteins annotated with the UniProt Keyword term ’Transmembrane’ were categorized as TM proteins. Among TM proteins, proteins annotated with the Gene Ontology Cellular Compartment term ’plasma membrane’ were classified as PM proteins, whereas others were classified as non-PM proteins. The solute carrier family members (SLCs) were classified as PM-associated SLCs (PM-SLCs) and non-PM-associated SLCs (non-PM-SLCs) as reported by Pizzagalli (Pizzagalli et al., 2021). The information about the topological domains and transmembrane regions of PM-proteins was obtained from UniProt.

The predicted structure of selected human PM-SLC and PM-associated TM (PM-TM) proteins was retrieved from AlphaFold Protein Structure Database (Green et al., 2022; Romera-Paredes et al., 2021). These included: SLC3A2 (P08195), SLC39A6 (Q13433), SLC6A8 (P48029), SLC7A5 (Q01650), SLC16A1 (P53985), SLC1A3 (P43003), CD44 (P16070), GPRC5A (Q8NFJ5), OSMR (Q99650), TGFBR1 (P36897), TFRC (P02786), EGFR (P00533), HLA-A (P04439), HLA-B (P01889), HLA-C (P43003), and BSG(Q54A51). The loosely folded regions of the AlphaFold-predicted structures were modified using Coot v0.9.4.1 (Emsley et al., 2010) to account for the spherical restriction of the plasma membrane with the preservation of the correct peptide bond conformation. The structures were visualized in Pymol (Schrödinger, v2.5.0) using cartoon representation. The protein’s extracellular, transmembrane, and extracellular regions were color-coded, and the lysine residues corresponding to the quantified UB sites were mapped.

##### Mapping high occupancy 40S sites in disome

To our knowledge, the structure of stalled human di-ribosome (disome) has not been reported. Thus, we used the structure of collided yeast disome (Ikeuchi et al., 2019) to map high occupancy 40S sites. The structure was retrieved from RCSB Protein Data Bank (PDB ID: 6I7O) and visualized in Pymol using a combination of cartoon and spherical representation. 40S proteins harboring the top one dozen high occupancy UB sites were represented in colors.

