## Supplemental Figures 1-7 for "A global, integrated view of the ubiquitylation site occupancy and dynamics"

### Figure S1

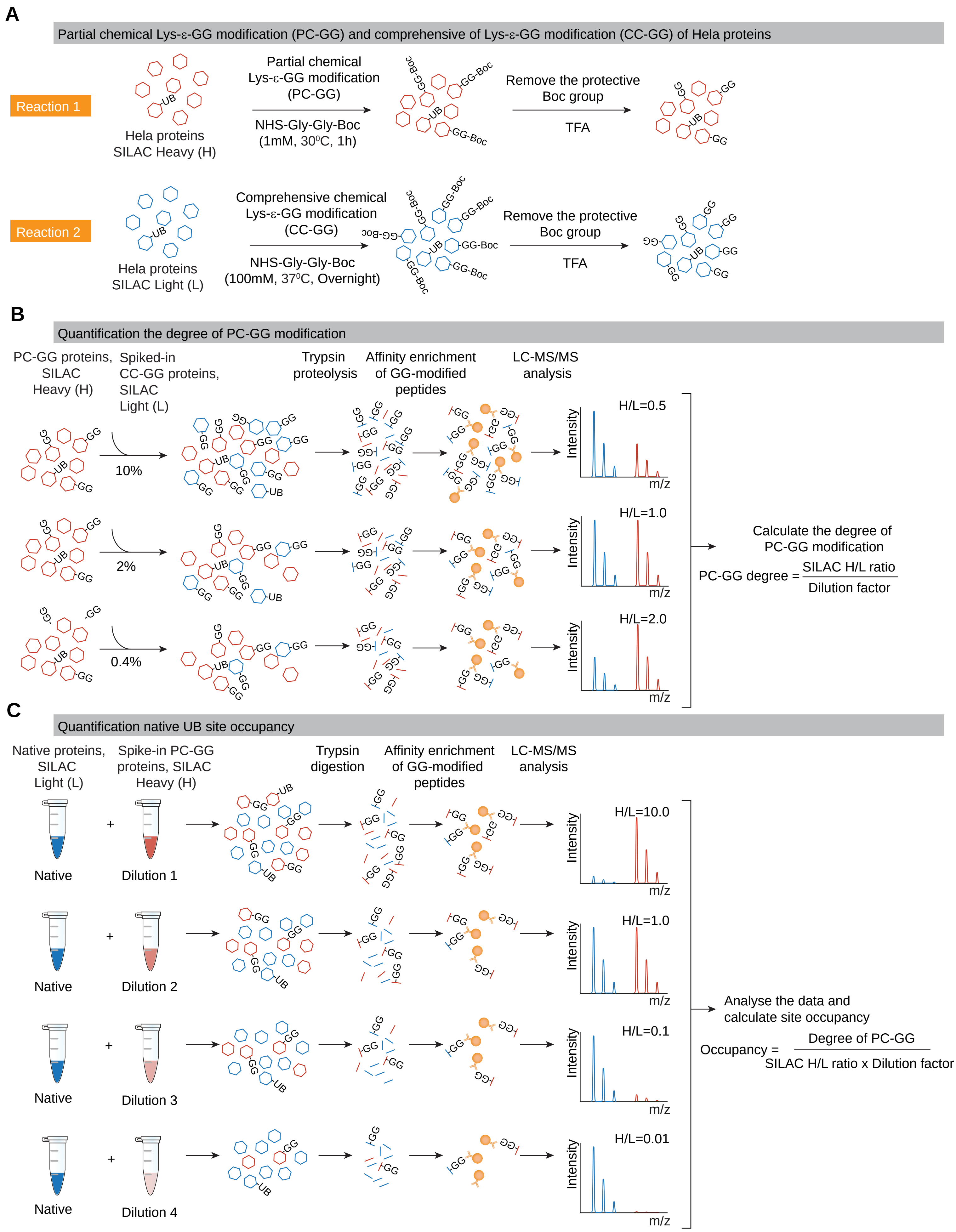

A
Partial chemical Lys-ε-GG modification (PC-GG) and comprehensive of Lys-ε-GG modification (CC-GG) of Hela proteins
Reaction 1
Hela proteins
SILAC Heavy (H)
Partial chemical
Lys-ε-GG modification
(PC-GG)
NHS-Gly-Gly-Boc
(1mM, 30⁰C, 1h)
Remove the protective
Boc group
TFA
Reaction 2
Hela proteins
SILAC Light (L)
Comprehensive chemical
Lys-ε-GG modification
(CC-GG)
NHS-Gly-Gly-Boc
(100mM, 37⁰C, Overnight)
Remove the protective
Boc group
TFA
B
Quantification the degree of PC-GG modification
PC-GG proteins, SILAC Heavy (H)
Spiked-in CC-GG proteins, SILAC Light (L)
Trypsin proteolysis
Affinity enrichment of GG-modified peptides
LC-MS/MS analysis
10%
2%
0.4%
H/L=0.5
H/L=1.0
H/L=2.0
Intensity
m/z
Calculate the degree of PC-GG modification
PC-GG degree = SILAC H/L ratio / Dilution factor
C
Quantification native UB site occupancy
Native proteins, SILAC Light (L)
Spike-in PC-GG proteins, SILAC Heavy (H)
Trypsin digestion
Affinity enrichment of GG-modified peptides
LC-MS/MS analysis
Native
Dilution 1
Dilution 2
Dilution 3
Dilution 4
H/L=10.0
H/L=1.0
H/L=0.1
H/L=0.01
Analyse the data and calculate site occupancy
Occupancy = Degree of PC-GG / SILAC H/L ratio x Dilution factor

**Figure S1**. **Strategy for the proteome-scale, site-specific quantification of ubiquitylation occupancy, related to Figure 1 and STAR Methods.** The schematic representation of the workflow for the site-specific quantification of ubiquitylation occupancy. For simplicity, the workflow is divided into three different steps (A-C). (**A**) Generation of partially chemically GG modified (PC-GG) protein standard and comprehensively chemically GG modified (CC-GG) protein standards. To generate PC-GG modified protein reference (reaction 1), SILAC heavy-labeled HeLa proteins were partially chemically modified using NHS-Gly-Gly-Boc and the indicated reaction conditions. To generate the CC-GG standard (reaction 2), SILAC-light-labeled HeLa proteins were modified using NHS-Gly-Gly-Boc and the indicated reaction conditions. The protective Boc protective group was removed using TFA (33%). (**B**) Quantification of the degree of PC-GG modification. To determine the degree of PC-GG modification, the CC-GG-modified proteins were mixed with PC-GG-modified proteins in different proportions. The mixed proteins were trypsin digested, GG-modified peptides were affinity-enriched, and analyzed by mass spectrometry (MS). The degree of PC-GG modification was calculated based on the SILAC ratio of GG-modified peptides from PC-GG and CC-GG samples, corrected for the dilution factor. (**C**) Quantification of the native UB site occupancy. To determine the occupancy of native UB sites, the reference PC-GG proteins were serially diluted and mixed with SILAC-light-labeled native HeLa proteins. The amount of spiked-in PC-GG protein was adjusted to yield the final site occupancy of 3% or 1%, 0.1%, 0.01%, and 0.001%. Subsequently, the mixed proteins were digested with trypsin, GG-modified peptides were affinity-enriched, and quantified by mass spectrometry. The native UB site occupancy is calculated using the SILAC ratio of native and chemically GG-modified peptides, the degree of PC-GG, and the PC-GG dilution factor, according to the indicated formula.

#### Figure S2

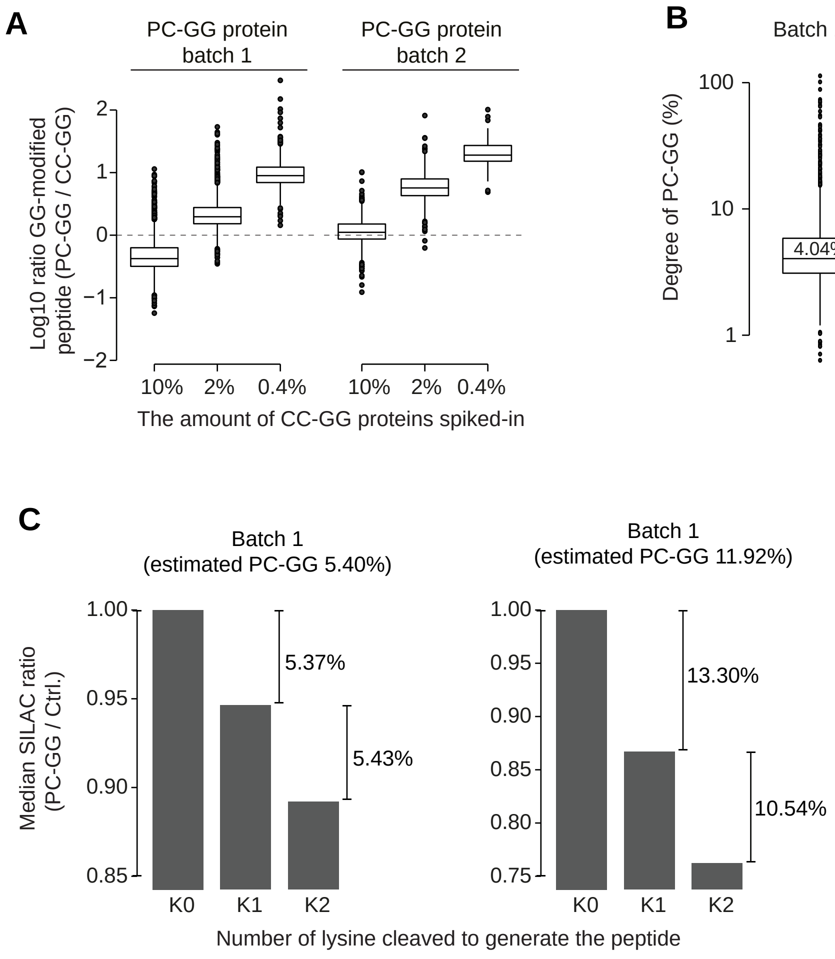

A
PC-GG protein batch 1
PC-GG protein batch 2
Log10 ratio GG-modified peptide (PC-GG / CC-GG)
2
1
0
−1
−2
10%
2%
0.4%
10%
2%
0.4%
The amount of CC-GG proteins spiked-in
B
Batch 1
Batch 2
Degree of PC-GG (%)
100
10
1
4.04%
11.28%
C
Batch 1
(estimated PC-GG 5.40%)
Batch 1
(estimated PC-GG 11.92%)
Median SILAC ratio (PC-GG / Ctrl.)
1.00
0.95
0.90
0.85
5.37%
5.43%
1.00
0.95
0.90
0.85
0.80
0.75
13.30%
10.54%
K0
K1
K2
K0
K1
K2
Number of lysine cleaved to generate the peptide

**Figure S2. Quantification of the degree of partial chemical GG modification, related to Figure 1 and STAR Methods.** (**A-B**) Quantification of the degree of partial chemical GG (PC-GG) modification by using comprehensively chemically GG (CC-GG) modified proteins. We prepared two independent batches (Batch 1 and Batch 2) of PC-GG-modified proteins. To quantify the degree of PC-GG modification, SILAC light-labeled CC-GG modified proteins were spiked-in into SILAC-heavy labeled PC-GG modified proteins in three different proportions, yielding the final CC-GG modification occupancy of 10%, 2%, and 0.4%. For each dilution, the box plot shows the SILAC ratio of GG-modified peptides in PC-GG and CC-GG samples, panel **A**. Using the data from panel **A**, the median degree of PC-GG modification was calculated with the indicated formula **B**. We only used peptides that were generated by trypsin cleavage at an arginine and following the dilution factor (with up to 2-fold variability). The peptides that were generated by trypsin cleavage at a lysine were excluded from these analyses. Box represents the interquartile range, the middle line denotes the median, and the whiskers indicate the minimum and maximum values, excluding outliers. (**C**) Estimating the degree of PC-GG modification by the measurement of lysine-dependent, unmodified peptides. Fully cleaved, unmodified peptides were classified based on the number and type of amino acids involved in tryptic digestion. The K0 peptides are generated by the cleavage at two arginines, the K1 peptides are generated by cleavage at one lysine and one arginine, and the K2 peptides are generated by cleavage at two lysines. Shown is the median SILAC ratio between PC-GG and untreated samples for each of the peptide types, normalized by the median SILAC ratio for K0 peptides. The difference between the K1-K0 peptides and K2-K1 peptides corresponds to the degree of PC-GG. This is because partial chemical GG modification prevents trypsin cleavage at the modified lysine, and the median decrease in the intensity of unmodified peptides generated by trypsin cleavage at lysine corresponds to the degree of PC-GG. K2 peptides are twice as likely to be impacted by the PC-GG modification than K1 peptides, and hence, the abundance of K2 peptides is decreased ~2-fold more than K1 peptides. K0 peptides are not expected to be impacted by the chemical modification, and thus, are used as a reference.

### Figure S3

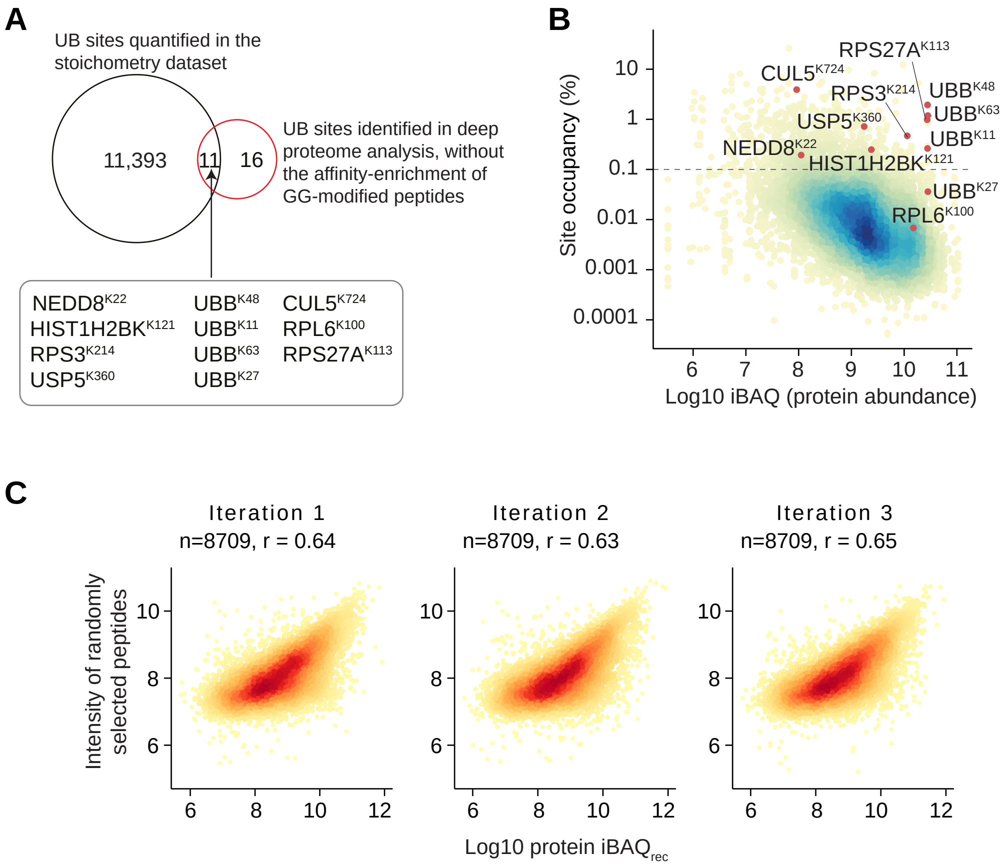
A
UB sites quantified in the stoichometry dataset
11,393
11
16
UB sites identified in deep proteome analysis, without the affinity-enrichment of GG-modified peptides
NEDD8K22
HIST1H2BKK121
RPS3K214
USP5K360
UBBK48
UBBK11
UBBK63
UBBK27
CUL5K724
RPL6K100
RPS27AK113
B
Site occupancy (%)
Log10 iBAQ (protein abundance)
RPS27AK113
CUL5K724
RPS3K214
UBBK48
UBBK63
USP5K360
UBBK11
NEDD8K22
HIST1H2BKK121
UBBK27
RPL6K100
C
Iteration 1
n=8709, r = 0.64
Iteration 2
n=8709, r = 0.63
Iteration 3
n=8709, r = 0.65
Intensity of randomly selected peptides
Log10 protein iBAQrec

**Figure S3. Validation of UB site occupancy quantification, related to Figure 1.** (**A**) The Venn diagram shows the overlap between UB sites identified in our UB site occupancy measurements (with affinity enrichment of GG-modified peptides) and the high confidence UB sites identified in a deep analysis of HeLa proteome, without the affinity-enrichment for GG-modified peptides (Bekker-Jensen et al., 2017). The total number of UB sites identified, and the sites identified in both datasets are specified. (**B**) The scatter plot shows the relationship between ubiquitylation occupancy calculated in this study and the iBAQ-based abundance of the corresponding proteins. The high confidence UB sites identified in deep total HeLa proteome analyses, without the affinity enrichment of GG-modified peptides, are indicated (red dots). iBAQ-based protein abundance was estimated using previously published HeLa cell proteome (Bekker-Jensen et al., 2017). (**C**) Estimation of the empirical correlation between MS intensity of individual peptides and the abundance of the corresponding proteins. Shown is the correlation between the intensity of a set of randomly selected unmodified peptides and the recalculated iBAQ abundance ($iBAQ_{rec}$) of the corresponding protein. For these analyses, we only chose proteins that were identified with at least 5 peptides. For each protein, 1 peptide was arbitrarily chosen to obtain a set of peptides with measured MS intensities, and the remaining peptides were used to recalculate the iBAQ-based abundance of the corresponding protein ($iBAQ_{rec}$). In this way, protein $iBAQ_{rec}$ was calculated after excluding the intensity of the randomly chosen peptide. To avoid any selection bias, we performed three iterations to select random peptides. The number of analyzed peptides (n) and Pearson's correlation (r) are shown.

#### Figure S4

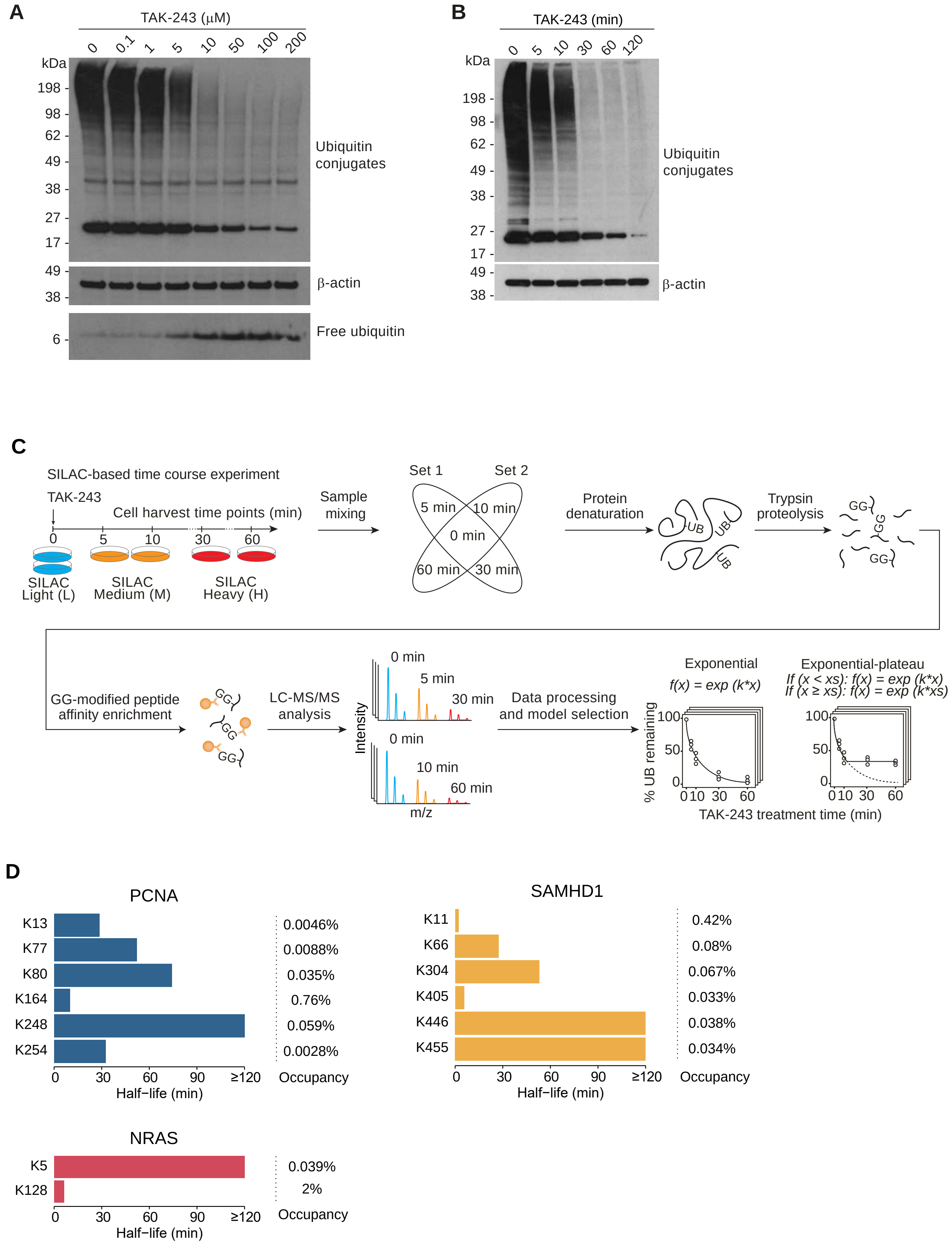

A
TAK-243 (µM)
0 0.1 1 5 10 50 100 200
kDa
198
98
62
49
38
27
17
Ubiquitin conjugates
49
38
β-actin
6
Free ubiquitin
B
TAK-243 (min)
0 5 10 30 60 120
kDa
198
98
62
49
38
27
17
Ubiquitin conjugates
49
38
β-actin
C
SILAC-based time course experiment
TAK-243
Cell harvest time points (min)
0 5 10 30 60
SILAC Light (L)
SILAC Medium (M)
SILAC Heavy (H)
Sample mixing
Set 1
Set 2
5 min
10 min
0 min
60 min
30 min
Protein denaturation
UB
Trypsin proteolysis
GG
GG-modified peptide affinity enrichment
LC-MS/MS analysis
Intensity
0 min
5 min
30 min
10 min
60 min
m/z
Data processing and model selection
Exponential
f(x) = exp (k*x)
Exponential-plateau
If (x < xs): f(x) = exp (k*x)
If (x ≥ xs): f(x) = exp (k*xs)
% UB remaining
100
50
0
0 10 30 60
TAK-243 treatment time (min)
D
PCNA
K13 0.0046%
K77 0.0088%
K80 0.035%
K164 0.76%
K248 0.059%
K254 0.0028%
0 30 60 90 ≥120
Half−life (min)
Occupancy
SAMHD1
K11 0.42%
K66 0.08%
K304 0.067%
K405 0.033%
K446 0.038%
K455 0.034%
0 30 60 90 ≥120
Half−life (min)
Occupancy
NRAS
K5 0.039%
K128 2%
0 30 60 90 ≥120
Half−life (min)
Occupancy

**Figure S4. Dose-response curve of E1 inhibition and the strategy for proteome-scale quantification of ubiquitylation site half-life, related to Figure 2.** (**A**) Shown is the dose-response of TAK-243 for global ubiquitylation inhibition. The immunoblot shows protein-conjugated and free ubiquitin levels after treatment (1h) with the indicated concentrations of TAK-243 (n= 3). Beta-actin was used as a loading control. Note that TAK-243-induced decrease in ubiquitylation saturates at ~100µM. (**B**) Time-course analysis of TAK-243-induced decrease in ubiquitylation. Cells were treated TAK-243 (100µM) for the indicated time and change in ubiquitylation was analyzed by immunoblotting (n=2). (**C**) Schematic representation of the SILAC-based time course experiment for quantifying UB site half-life. Cells were encoded with triple SILAC (light, medium, and heavy), treated with TAK-243 (100µM) as specified, and lysed at the indicated time points after the treatment. To cover 5 different time points (0, 5, 10, 30, 60 minutes), two sets of SILAC experiments were performed. In set 1, proteins from cells treated for 0, 5, and 30 minutes were combined, and in set 2, proteins from cells treated for 0, 10, and 60 minutes were combined. For each set, the combined proteins were digested with trypsin, and GG-modified peptides were affinity-enriched and analyzed on a mass spectrometer. We fit two decay models: an exponential and exponential-plateau function to the kinetic profile. The better fitting model was selected based on Akaike's information criteria, and the UB site half-life is calculated based on the exponential part of the decay function. The experiment was performed in 5 biological replicates. (**D**) Ubiquitylation occupancy and half-life are controlled site-specifically. Representative examples of proteins showing site-specific differences in the half-life and occupancy. In PCNA, K164 has the shortest half-life and K248 has the longest half-life. In NRAS K128 has a fast half-life and K5 has a very long half-life. In SAMHD1, K11 has the fastest half-life and K445/K446 have the longest half-life. In these proteins, sites with the fastest turnover also have the highest occupancy, showing that the high occupancy of these sites is not due to their slow turnover but due to their targeted ubiquitylation.

**Figure S5**

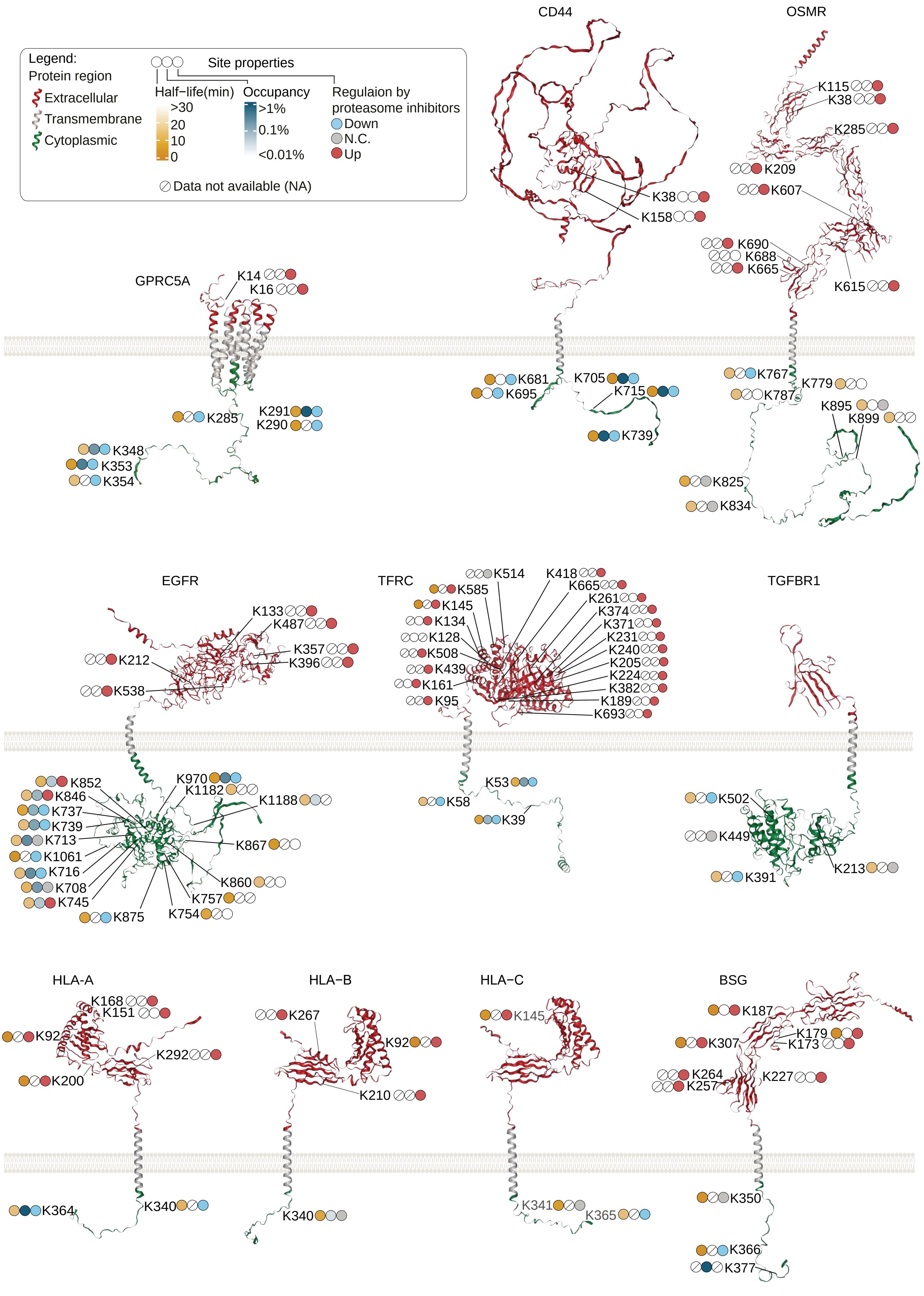

Legend:
Protein region
Extracellular
Transmembrane
Cytoplasmic
Site properties
Half−life(min)
>30
20
10
0
Occupancy
>1%
0.1%
<0.01%
Regulaion by proteasome inhibitors
Down
N.C.
Up
Data not available (NA)
CD44
K38
K158
K681
K695
K705
K715
K739
OSMR
K115
K38
K285
K209
K607
K690
K688
K665
K615
K767
K787
K779
K895
K899
K825
K834
GPRC5A
K14
K16
K285
K291
K290
K348
K353
K354
EGFR
K133
K487
K212
K357
K396
K538
K852
K846
K737
K739
K713
K1061
K716
K708
K745
K875
K970
K1182
K1188
K867
K860
K757
K754
TFRC
K514
K418
K585
K665
K145
K261
K374
K134
K371
K128
K231
K508
K240
K439
K205
K161
K224
K382
K95
K189
K693
K53
K58
K39
TGFBR1
K502
K449
K391
K213
HLA-A
K168
K151
K92
K292
K200
K364
K340
HLA-B
K267
K92
K210
K340
HLA-C
K145
K341
K365
BSG
K187
K179
K307
K173
K264
K257
K227
K350
K366
K377

**Figure S5. Distinct properties of the sites occurring in the intracellular and extracellular domains of the plasma membrane transmembrane receptors, related to Figure 4.** Shown are the properties of UB sites occurring in the indicated plasma membrane transmembrane proteins. UB sites were mapped to protein structures predicted by AlphaFold (Green et al., 2022; Romera-Paredes et al., 2021). Different topological domains (cytoplasmic, extracellular, transmembrane) are color-coded as indicated. In each protein, the identified UB sites, their occupancy, half-life, and regulation by proteasome inhibition are shown.

### Figure S6

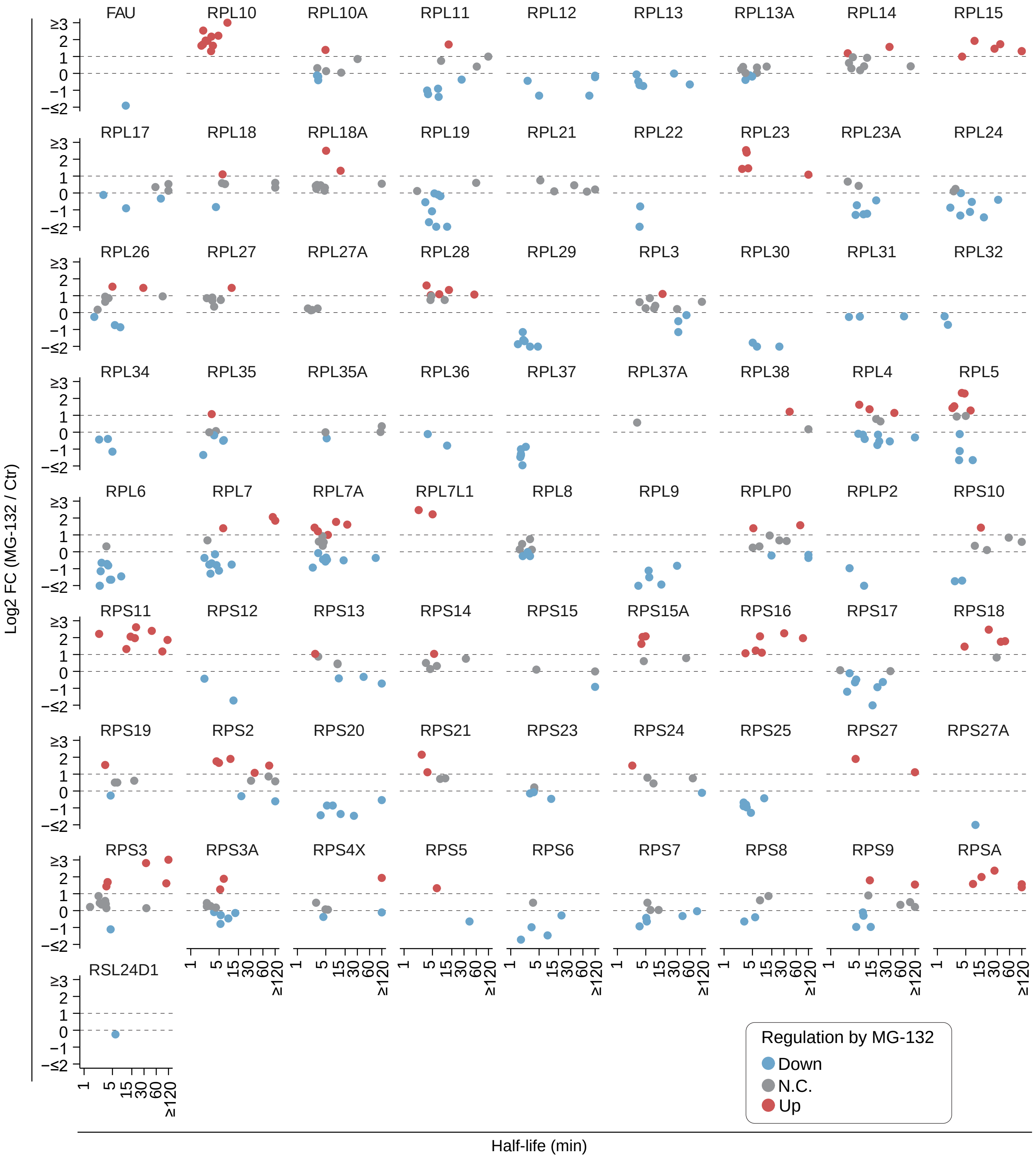

FAU RPL10 RPL10A RPL11 RPL12 RPL13 RPL13A RPL14 RPL15
RPL17 RPL18 RPL18A RPL19 RPL21 RPL22 RPL23 RPL23A RPL24
RPL26 RPL27 RPL27A RPL28 RPL29 RPL3 RPL30 RPL31 RPL32
RPL34 RPL35 RPL35A RPL36 RPL37 RPL37A RPL38 RPL4 RPL5
RPL6 RPL7 RPL7A RPL7L1 RPL8 RPL9 RPLP0 RPLP2 RPS10
RPS11 RPS12 RPS13 RPS14 RPS15 RPS15A RPS16 RPS17 RPS18
RPS19 RPS2 RPS20 RPS21 RPS23 RPS24 RPS25 RPS27 RPS27A
RPS3 RPS3A RPS4X RPS5 RPS6 RPS7 RPS8 RPS9 RPSA
RSL24D1
Log2 FC (MG-132 / Ctr)
≥3 2 1 0 −1 −≤2
1 5 15 30 60 ≥120
Half-life (min)
Regulation by MG-132
Down
N.C.
Up

**Figure S6. Ubiquitylation of the ribosome is regulated in a site- and protein-specific manner, related to Figure 5.** Shown is the half-life and regulation by MG-132 (2.5 h) of the sites occurring in the indicated ribosomal subunits. Notably, UB sites in many subunits, including RPL29, RPL37, RPL5, and RPL10, have similarly short half-lives, but they show distinct regulation upon MG-132 treatment.

#### Figure S7

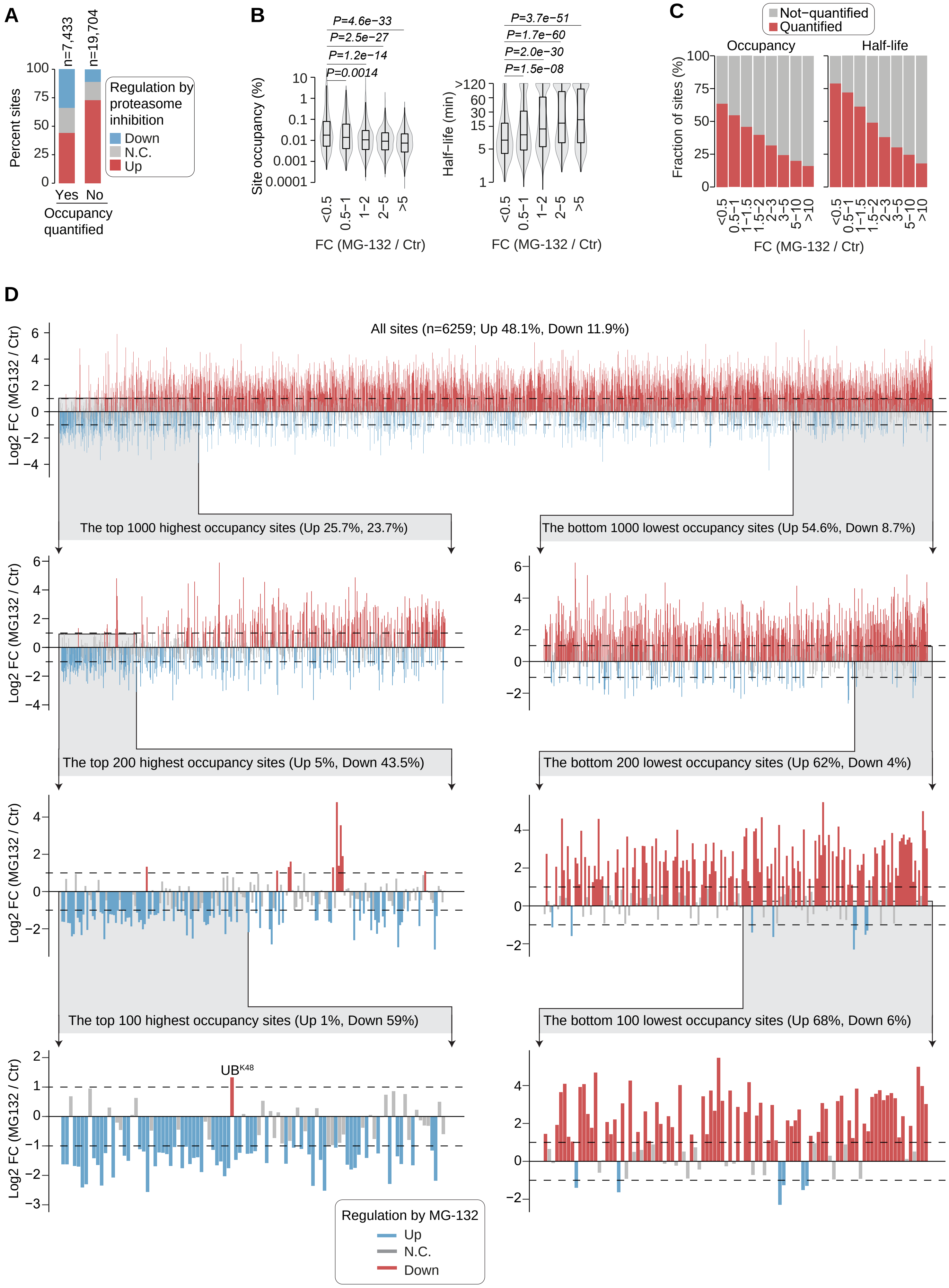
A
n=7,433
n=19,704
Percent sites
100
75
50
25
0
Yes No
Occupancy quantified
Regulation by proteasome inhibition
Down
N.C.
Up
B
P=4.6e−33
P=2.5e−27
P=1.2e−14
P=0.0014
Site occupancy (%)
FC (MG-132 / Ctr)
P=3.7e−51
P=1.7e−60
P=2.0e−30
P=1.5e−08
Half−life (min)
FC (MG-132 / Ctr)
C
Not−quantified
Quantified
Occupancy
Half-life
Fraction of sites (%)
FC (MG-132 / Ctr)
D
All sites (n=6259; Up 48.1%, Down 11.9%)
Log2 FC (MG132 / Ctr)
The top 1000 highest occupancy sites (Up 25.7%, 23.7%)
The bottom 1000 lowest occupancy sites (Up 54.6%, Down 8.7%)
The top 200 highest occupancy sites (Up 5%, Down 43.5%)
The bottom 200 lowest occupancy sites (Up 62%, Down 4%)
The top 100 highest occupancy sites (Up 1%, Down 59%)
The bottom 100 lowest occupancy sites (Up 68%, Down 6%)
UBK48
Regulation by MG-132
Up
N.C.
Down

**Figure S7. High and low occupancy sites are differentially regulated by proteasome inhibition, related to Figure 7.** (**A**) Shown is the fraction of UB sites quantified after MG-132 treatment (2.5h). Sites quantified in MG-132 treated cells were grouped based on whether they were quantified (Yes) or not (No) in our occupancy measurements. Within each group, the fraction of MG-132 regulated sites is shown. The number of UB sites (n) analyzed is donated. (**B**) Relationship between UB site occupancy (left panel) or half-life (right panel) with the change in ubiquitylation after MG-132 treatment (2.5h). Based on MG-132-induced fold-change in ubiquitylation, the sites were grouped into five different groups, and the groups were ordered with lowest to highest increase in ubiquitylation after MG-132 treatment. Within each group, site occupancy (left panel), and half-life (right panel) are shown. (**C**) With increasing upregulation by MG-132, the fraction of sites with quantified occupancy and half-life decreases. Based on MG-132-induced fold-change in ubiquitylation, the sites were grouped into eight groups, and the groups were ordered with lowest to highest increase in ubiquitylation after MG-132 treatment. Within each group, the fraction of sites with or without quantified occupancy (left panel), and half-life (right panel) are shown. (**D**) Shown is the distribution of UB site regulation by proteasome inhibition by MG-132 (2.5h). All UB sites with valid occupancy values, and quantified in response to MG-132, were ranked based on their occupancy (left-to-right, highest-to-lowest occupancy). The figures show all UB sites quantified, as well as the top and bottom-ranked 1000 sites, top and bottom-ranked 200 sites, and the top and bottom-ranked 100 sites. Each bar represents an individual UB site and its regulation after MG-132 treatment. Of note, among the top 100 highest occupancy sites, only one site is upregulated by MG-132, and this site, corresponding to K48 in ubiquitin ($UB^{K48}$), is marked.
