## Supplemental Notes 1-5 for "A global, integrated view of the ubiquitylation site occupancy and dynamics"

**Supplementary Note 1-5**

**Supplemental Note 1**

In HeLa, the abundance of MS-detectable proteins ranges from $<10^2$ to $>10^7$ copies/cell, with the estimated median protein abundance of ~$10^4$ copies/cell (Bekker-Jensen et al., 2017). 44% of the sites quantified in our occupancy dataset occurred in highly abundant proteins (with an estimated abundance of $>10^6$ copies/cell). If the sites occurring in these highly abundant proteins were ubiquitylated at >1% occupancy, the estimated abundance ($>10^4$ copies/cell) of ubiquitylated peptides would be in the same range as the median abundance of HeLa protein. Such high abundance should enable the detection of ubiquitylated peptides without their affinity enrichment. To test this idea, we searched a deep HeLa cell proteome dataset for GG-modified peptides (Bekker-Jensen et al., 2017). As a reference, the same dataset was simultaneously searched for phosphorylation, which is known to occur at a broad range of occupancies (Olsen et al., 2010; Wu et al., 2011). A standard database search (at a false discovery rate of 1%), identified 9,690 phosphorylated peptides but only 350 GG-modified peptides (**Figure 1D**). Searching MS data with multiple variable modifications increases the database search space and leads to a higher rate of false positives among peptides that are identified with low abundant, infrequently occurring PTMs (Bogdanow et al., 2016). To eliminate low-confidence identifications, we applied stringent filtering criteria and removed peptides bearing multiple variable modifications and low identification scores (**see Methods**). This resulted in the removal of a disproportionately high (90%) fraction of GG-modified peptides as compared to unmodified and phosphorylated (15-17%) peptides (**Figure 1D**). After filtering, we detected 309,053 unmodified, 8,036 phosphorylated, and 35 GG-modified peptides, harboring 27 unique UB sites (**Figure 1D**). Of these UB sites, 11 overlapped with the sites quantified in our occupancy dataset (**Figure S3A**). The overlapping sites had high occupancy and/or occurred in highly abundant proteins (**Figure S3B**). These results show that, without their affinity-enrichment, the chances of detecting GG-modified peptides in MS-based proteome analyses are ~230 times lower than phosphorylated peptides and ~8,800 times lower than unmodified peptides. This supports low occupancy of ubiquitylation and explains why ubiquitylated peptides are seldom detected without their affinity enrichment.

**Supplemental Note 2**

Because of the differences in peptide ionization efficiencies, and other unknown factors, the MS intensity of peptides does not always scale directly with their abundance. Nonetheless, MS intensity generally correlates with peptide abundance, and MS intensity is widely used for estimating absolute protein abundance using label-free methods, such as Top3 and iBAQ (Ahrne et al., 2013; Schwanhäusser et al., 2011; Silva et al., 2006). For posttranslationally modified peptides, the abundance of peptides reflects the abundance of protein from which it is derived, as well as the site occupancy of the PTM. Previously, we showed that protein abundance-corrected intensity of acetylated peptides (Weinert et al., 2015), which we refer to as abundance-corrected modified peptide intensity (ACI), correlates with the modification site occupancy.

To validate our UB site occupancy estimates, we determined ACI for GG-modified peptides by normalizing the GG-modified peptide intensity by the abundance of corresponding proteins. Protein abundance was estimated using iBAQ (Schwanhäusser et al., 2011). ACI of GG-modified peptides shows a notable correlation with iBAQ (r= 0.70) (**Figure 1E**). Because peptide intensity does not strictly correlate with the abundance, the ideal correlation between the ACI of GG-modified peptides and protein iBAQ is not expected to be 1. How well does the peptide intensity correlate with iBAQ-based protein abundance? To address this question, we looked at the correlation between the intensity of individual peptides and the iBAQ-based abundance of the corresponding protein. The idea is that proteolysis of a protein typically generates multiple peptides, which should be present in equimolar amounts (assuming that the protein is not modified with PTMs, or the abundance of PTM is negligible). Therefore, the intensity of individual peptides and the iBAQ-based corresponding protein abundance should reflect the empirical correlation. To determine the empirical correlation, we used HeLa proteins identified with ≥5 peptides, and from each protein, we randomly chose one peptide.

Protein iBAQ is calculated as defined previously (Schwanhüusser et al., 2011).

$$iBAQ = \frac{Int_{all}}{n_{theor}}$$

where: $iBAQ$ - protein iBAQ value, $Int_{all}$ - the summed intensity of all peptides, $n_{theor}$- number of theoretically observable tryptic peptides.

To eliminate the contribution of the intensity of the randomly chosen peptide to the calculation of protein abundance, the corresponding protein iBAQ was re-calculated as follows:

$$iBAQ_{rec} = \frac{Int_{all} - Int_{random}}{n_{theor} - 1}$$

where: $iBAQ_{rec}$- recalculated protein iBAQ value, $Int_{all}$ - the summed intensity of all peptides, $Int_{random}$ - the intensity of the randomly selected peptide, $n_{theor}$- number of theoretically observable tryptic peptides.

To rule out a bias in the random peptide selection, we performed three iterations by choosing a different set of random peptides. Overall, the intensity of individual peptides showed a good correlation (r=0.63-65) with the abundance of the corresponding protein (**Figure S3C**). Notably, the correlation between UB site occupancy and the ACI of GG-modified peptides is even higher (r=0.70) than expected maximum correlation (r=0.63-65) (**Figure S3C**). This is possible because GG-modified peptides carry an extra positive charge (due to an additional amino group from the GG remnant), which likely improve their ionization efficiency. The concordance between ACI and UB site occupancy estimates is striking when considering all the possible sources of variability in our analyses, including inherent differences in peptide ionization, site-specific variation in chemical modification, variation in antibody enrichment of GG-modified peptides, and quantification errors in different MS measurements. Overall, the excellent correlation between experimentally measured UB site occupancy and the ACI of the GG-modified peptides, which are measured entirely independently of each other, supports the accuracy of PC-GG-based occupancy estimates.

**Supplemental Note 3**

If we know the total number of protein molecules, the number of ubiquitin molecules, the fraction of ubiquitin molecules conjugated to proteins, and the number of ubiquitylatable lysine residues in a cell, it should be possible to obtain a theoretical estimate of the global UB site occupancy. To obtain the required parameters, we performed experimental measurements and used known values from published literature. Using a synthetic peptide reference, we estimated a ubiquitin concentration of 477±68 pmol/mg of HeLa proteins, which is in good agreement with the previously reported ubiquitin concentration in HEK293 cells (486.4 ± 42 pmol/mg) (Kaiser et al., 2011). Total protein concentration in HeLa cells was estimated at 158±15 pg/cell, which is also consistent with the prior estimates (150 pg/cell) (Volpe and Eremenko-Volpe, 1970). By using the estimated total protein and ubiquitin content per cell, we estimated $4.54 \times 10^7$ ubiquitin molecules/cell in HeLa. A HeLa cell contains ~$4 \times 10^9$ protein molecules/cell (Bekker-Jensen et al., 2017), therefore, we estimate that ubiquitin constitutes ~1.1% of the total protein molecules. If all of the ubiquitin was conjugated to proteins, and every protein was modified on just 1 lysine, the maximum average occupancy would be 1.1%. In cultured human and mouse cells, 77% of ubiquitin is conjugated to proteins and the remaining ubiquitin is present as a free pool (Kaiser et al., 2011). If 77% of the total ubiquitin pool was conjugated to proteins, it can modify 0.85% of protein molecules in HeLa, on a single lysine. Most proteins are not just modified at a single lysine but at multiple lysine residues. Cumulatively, MS analyses have identified >100,000 UB sites, and up to ~90,000 sites have been identified in a single study in a single cell type (Hansen et al., 2021; Kim et al., 2011; Pinto-Fernandez et al., 2020; Steger et al., 2021; Wagner et al., 2011; Wilson et al., 2018). From the number of experimentally identified UB sites, it can be conservatively estimated that an average human protein is ubiquitylated at ~10 lysines. Because the number of ubiquitylatable lysines is ten times greater than the number of protein molecules, the average theoretical UB site occupancy will be 0.085%. The distribution of protein abundance is skewed, and in HeLa, the median protein abundance is more than an order of magnitude lower than the average abundance (Bekker-Jensen et al., 2017). If ubiquitylation occupancy follows the same distribution pattern, the theoretical median and mean UB site occupancy will be 0.0085% and 0.085%, respectively. These theoretical estimates are in good agreement with our experimentally measured median (0.0081%) and mean (0.059%) occupancy of ubiquitylation.

**Supplementary Note 4**

To our knowledge, only one study has used MS to quantify the site-specific half-life of PTMs on a large scale (i.e., >100 sites), in living cells (Zee et al., 2010). To measure the turnover rate of lysine acetylation, cells were cultured in media containing $^{13}C$-glucose, which is metabolized into acetyl-CoA, which serves as an essential precursor for acetylation (Zee et al., 2010). Acetylation turnover rates were determined by quantifying the rate of $^{13}C$ incorporation into acetylated peptides. By quantifying ~900 sites, the median half-life of acetylation was estimated to be ~33h (Kori et al., 2017), which is only slightly lower than the median half-life of proteins in HeLa (37h) (Cambridge et al., 2011; Zecha et al., 2018). A major caveat of this approach is the slow conversion of $^{13}C$-glucose to acetyl-CoA; it takes ~8h before the level of $^{13}C$-labeled acetyl-CoA equalizes with the $^{12}C$-containing acetyl-CoA. The slow conversion of glucose to acetyl-CoA and co-existence of the $^{13}C$ and $^{12}C$ labeled acetyl-CoA pools make accurate quantification difficult and lead to a severe underestimation of the half-life. Indeed, using a selective chemical inhibitor of CBP/p300, we quantified the half-lives ~600 sites and the median half-life was estimated to be ~1h (Weinert et al., 2018). However, a lack of broad-range acetyltransferase inhibitors prevented global measurement of acetylation site half-life.

**Supplemental Note 5**

To achieve rapid and complete inhibition of E1 and the proteasome, we used fully saturating concentrations of the chemical inhibitors. At these high concentrations, the inhibitors may exhibit some off-target effects, for example, a high concentration of TAK-243 can inhibit enzymes involved in regulating NEDD8 or ISG15 (Hyer et al., 2018). However, the use of sub-saturating inhibitor concentrations risks underestimating the true kinetics of ubiquitylation. Also, the GG profiling approach cannot anyway discriminate UB sites from sites modified by NEDD8 and ISG15, which constitute a small fraction (<5%) of GG-modified peptides (Kim et al., 2011).
